# A new polymodal gating model of the proton-activated chloride channel

**DOI:** 10.1101/2022.10.31.514568

**Authors:** Piao Zhao, Cheng Tang, Yuqin Yang, Zhen Xiao, Samantha Perez-Miller, Heng Zhang, Guoqing Luo, Hao Liu, Yaqi Li, Qingyi Liao, Fan Yang, Hao Dong, Rajesh Khanna, Zhonghua Liu

**Author notes:** Correspondence to: C.T.; H.D.; R.K. and Z.L. These authors contributed equally to this work (Piao Zhao, Cheng Tang, Yuqin Yang).

## Abstract

The proton-activated chloride (PAC) channel plays critical roles in ischemic neuron death, but its activation mechanisms remain elusive. Here, we interrogated PAC channel gating using its unique bidirectional modulator C77304 as a pharmacological probe. C77304 activated the PAC channel by acting on its proton gating, while simultaneously inhibiting channel activity at higher doses, through interaction with two modulatory sites with different affinities and state-dependence. Excitingly, we revealed that PAC undergoes intrinsic proton gating-independent voltage activation, which was defined by an ion-flux gating mechanism. Scanning-mutagenesis and molecular dynamics simulation confirmed that E181, E257, and E261 in human PAC form the primary proton sensors, as alanine mutations eliminated the channel’s proton gating while sparing the voltage-dependent gating. This proton sensing mechanism was basically conserved among orthologous PAC channels. Collectively, our data unveils the polymodal gating and proton sensing mechanisms in the PAC channel which may inspire potential drug development.

## Introduction

Proton, possibly the smallest ligand in biology, regulate the activity of various ion channels on plasma and intracellular membranes by modifying the channels’ response to other gating stimuli^1-4^, or by acting as direct channel agonist^5^, or both^6^. While the proton-activated, acid-sensing cation channels (ASICs) were identified decades ago and have been intensively studied^7^, the proton activated chloride (PAC) channel contributing to acid-sensing, outwardly rectifying (ASOR) anion currents, was only recently characterized^8-10^. The PAC channel is a homotrimer, with each subunit composed of two transmembrane segments (TM1, TM2) connected by a spanning extracellular loop (ECL)^8,9,11-13^. Spatially, the pore-lining helix TM2 and the peripheral TM1 create the pore domain (PD), while the ECL forms the extracellular domain (ECD); the ECD constricts at the extracellular membrane face to join the PD^11-13^. A critical lysine residue at the intracellular end of TM2 serves as the Cl^-^ selectivity filter^11^. Both the topological and spatial structures of PAC bear similarity to ASICs and the ATP-gated purinergic P2X receptor cation channels^14,15^. The large, hand-like ECD domain in PAC likely contains multiple sites for pharmacological modulation, as that in ASICs^16^. However, unlike ASICs, the ECD of PAC lacks the exterior helix domain which comprises the presumptive acid pocket, suggesting that it may work via a specialized proton sensing mechanism^11^.

Structural biology approaches have offered insights into possible mechanisms of proton sensing in PAC. Comparison of the resting and desensitized PAC structures implicated a histidine residue at the extracellular end of TM1 (H98 – human PAC numbering) as the proton sensor, with S102, Q296 and I298 forming its docking site (pH8.0); in the protonated state (pH4.0), the sensor is believed to reside in the “ acid pocket” formed by E107, D109 and E250^11^. However, mutating these residues did not attenuate the channel’s proton sensitivity, raising questions as to their roles in proton sensing. Translating the differences between the resting and the putative desensitized, but not the true opened, PAC structures to conformational changes initiated by acid sensing might be problematic. Comparing the resting and activated PAC structures, the Long group proposed another proton sensing mechanism, in which six titratable residues likely create the proton sensor and form pairwise interactions in the channel’s activated state (E257-D289, E249-E107, and E250-D297)^13^. However, it’s unknown whether such interactions are the initial driving force for proton gating. Consequently, the proton sensing mechanism in PAC remains unknown, as that in ASICs^17^. The gating mechanism of PAC could be much more complicated than that of ASICs, for which a depolarization-facilitated gating was reported^18^. Since PAC does not possess a prototypical voltage sensor commonly found in classic voltage-gated ion channels^19^, they may sense voltage via an unorthodox mechanism.

As an evolutionarily conserved protein^8,9^, the PAC channel plays critical physiological and pathological roles. Reducing or abolishing PAC activity protects cells from acid-induced necrotic cell death in vitro and in animal models^8,20,21^. PAC has also been reported to function as pH sensors to prevent hyper-acidosis in endosomes^22^. Therefore, pharmacological agents modulating PAC activity can be leveraged as molecular probes to investigate key structure-function relationships. Unfortunately, only non-specific PAC antagonists abound, and their mechanism of action remains unknown^21,23-25^.

Herein, we explored PAC channel gating using its novel and unique bidirectional modulator C77304 (5-iodo-2-(2-methylfuran-3-carboxamido)benzoicacid) as a pharmacological tool. By uncovering C77304’s action mechanism, we demonstrated, for the first time, that the PAC channel undergoes proton gating-independent voltage gating. Scanning-mutation analysis and molecular dynamics simulations revealed that the E181, E257, and E261 mutations in human PAC specifically abolished the channel’s proton gating but spared its voltage-dependent gating. Cross-species analysis confirmed that this proton sensing mechanism is both conserved and variable among orthologous PAC channels. Altogether, our data advances foundational knowledge of the gating mechanism of PAC channel, which may spur interest in targeting it for drug development.

## Results

### C77304 bidirectionally modulates PAC activity

Pharmacological agents interfering with the proton gating of the PAC channel are valuable molecular tools for dissecting its gating mechanism. We screened a compound library for such agents and found a hit molecule (C77304; upper panel in Fig. 1A, Supplementary Figs. 1-2) that concentration-dependently inhibited PAC currents at pH4.6 (Fig. 1B). Increasing the proportion of resting-state channels by recording at pH5.34 produced a bimodal response of C77304: at 10 µM, PAC currents were potentiated by ∼2-3 fold (Fig. 1C), while at 50 µM, the currents were robustly inhibited (Fig. 1D). The activation effect was not caused by pH fluctuation by C77304 dissociation [lower panel in Fig. 1A]. These data suggest that C77304’s effects might be driven by a competition between its activation and inhibition actions. To probe this further, we performed time-course experiments with C77304. PAC currents (at pH5.6 - most channels are in the resting state) were transiently activated by acute application of 50 µM C77304 but then were gradually inhibited (Fig. 1E). However, pretreatment with 1 µM C77304, to favor PAC channel opening, eliminated transient activation and now the currents were directly inhibited by C77304 (Fig. 1F). The concentration-response curves of C77304 against the PAC channel were bell-shaped in relatively weak acidic conditions (i.e., pH5.34, pH5.6, pH5.8), demonstrating bidirectional modulation (Fig. 1G). Contrarily, C77304 monotonically inhibited PAC currents at pHs below 5.0 due to the overwhelming inhibitory effect (Fig. 1G). The divergence between the activation and inhibition phenotypes is likely because the EC_50_ for C77304 activating the PAC channel was roughly 9–25 fold lower than its IC_50_ in blocking it (Figs. 1H-I). The inhibitory effect of C77304 on PAC was slightly attenuated in conditions where a bidirectional modulation was observed (Fig. 1H), suggesting that the compound inhibited the proton-activated and the proton- and compound-activated channels with different potencies and/or efficacies. Testing the binding and unbinding kinetics of C77304 on the PAC channel showed that the time-course for current activation was faster than that of inhibition; both were reversible (Figs. 1J-K). These data strongly imply simultaneous binding of C77304 to the activation and inhibition sites on the channel with differing affinities (see below).

**Fig 1.**
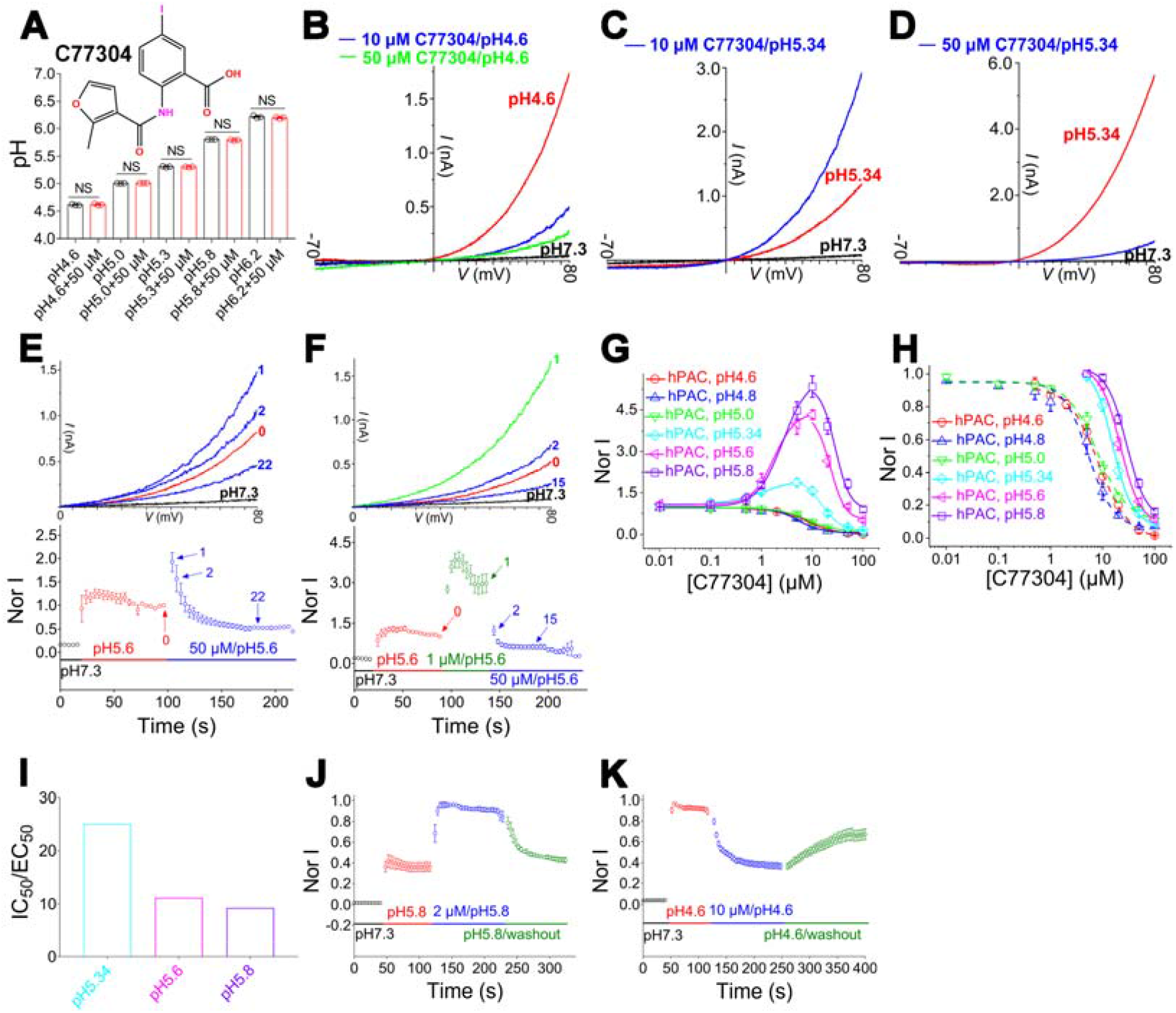
Identification of C77304 as a bidirectional modulator of PAC channel. (A), upper panel, C77304 structure; lower panel, C77304 does not change the buffer pHs (n = 3; NS, not significant; paired t-test). (B), C77304 concentration-dependently inhibits PAC currents elicited by ramp depolarizations from -70 mV to +80 mV at pH4.6 (n = 6). (C)-(D), C77304 at 10 µM potentiates (C), but at 50 µM inhibits (D), PAC currents at pH5.34 (n = 15). (E)-(F), Time course of C77304 acting on PAC channels at pH5.6. Currents were normalized to the current labeled as 0. Inset: example current traces at different time points as indicated (n = 5-7). (G), concentration-response curves of C77304 activating and/or inhibiting PAC currents at different pHs. The activation EC_50_s and mean hill slopes were 2.9 ± 0.9 μM and 1.6 (n = 19), 1.8 ± 0.6 μM and 1.9 (n = 8), and 0.6 ± 0.2 μM and 2.1 (n = 16) at pH5.8, pH5.6, and pH5.34, respectively. (H), concentration-response curve of C77304 inhibiting PAC currents at different pHs, currents were normalized to their respective maximum compound-activated currents at pH5.8, pH5.6, and pH5.34. The IC_50_s and the mean hill slopes were 26.9 ± 2.6 μM and 1.9 (n = 19), 20.2 ± 1.5 μM and 2.5 (n = 8), 15.5 ± 1.1 μM and 2.6 (n = 15), 7.9 ± 0.5 μM and 2.0 (n = 8), 6.2 ± 0.8 μM and 1.8 (n = 7), and 8.3 ± 1.0 μM and 1.5 (n = 6) at pH5.8, pH5.6, pH5.34, pH5.0, pH4.8, and pH4.6, respectively. (I), The ratio of the apparent affinity of C77304 inhibiting and activating PAC channels (IC_50_/EC_50_) at pHs as indicated. (J)-(K), Time-course of C77304 activating (J) and inhibiting (K) PAC currents and the corresponding wash-out upon perfusion with compound-free solutions. The calculated association time constant (τ_on_) was 4.5 ± 1.3 s and 15.9 ± 2.3 s, and the dissociation time constant (τ_off_) was 15.9 ± 2.5 s and 82.7 ± 12.9 s, at pH5.8 and pH4.6, respectively (n = 6-10).

### A model for C77304 bidirectionally modulating the PAC channel

The interaction sites on PAC channel for C77304’s activation, inhibition, and proton binding were defined as sites (1), (2) and (0), respectively. Thus, there exist eight combinations of their occupation states. Theoretically, there are simultaneously sixteen different channel states depending on whether the channel’s pore is open or closed. We simplified the model to contain eight practically achievable states based on experimental data (Fig. 2A). The state-dependence of C77304 binding to site 2 was analyzed using the experimental protocol (Fig. 2B; upper panel), in which currents were evoked by two pH5.0 challenges separated by pH7.3 preincubation with or without compound. A saturating concentration (100 µM) of C77304 preincubated with wild-type (wt) PAC channel at pH7.3 did not inhibit the currents (Figs. 2B-C), suggesting that site 2 was not accessible to C77304 in the channel’s resting state. In contrast, the A321C mutant channel, which showed basal opening at pH7.3^9^, was inhibited with this preincubation treatment (Figs. 2B-C). Preincubation of both channels with bath solution (pH7.3 only) did not affect their currents (Figs. 2B-C). The kinetics of preincubated C77304 inhibiting PAC currents also demonstrated that the inhibition was accompanied by and lagged channel activation. A 5-min preincubation with 100 μM C77304 at pH7.3 was expected to fully inhibit PAC currents if the compound binds to the resting channels and accordingly a subsequent pH4.6 test would detect no channel activity. However, as shown in Fig. 2D, ∼20% of PAC currents remained at the beginning of the pH4.6 pulse, which was then quickly inhibited as the channels became activated. Most (∼80%) channels might be immediately inhibited post activation, due to close proximity to C77304 and the fast inhibition kinetics. Collectively, these data demonstrate that C77304 inhibits the PAC channel by promoting O_1_→C_4_ and O_2_→C_5_ state transitions (Fig. 2A).

**Fig 2.**
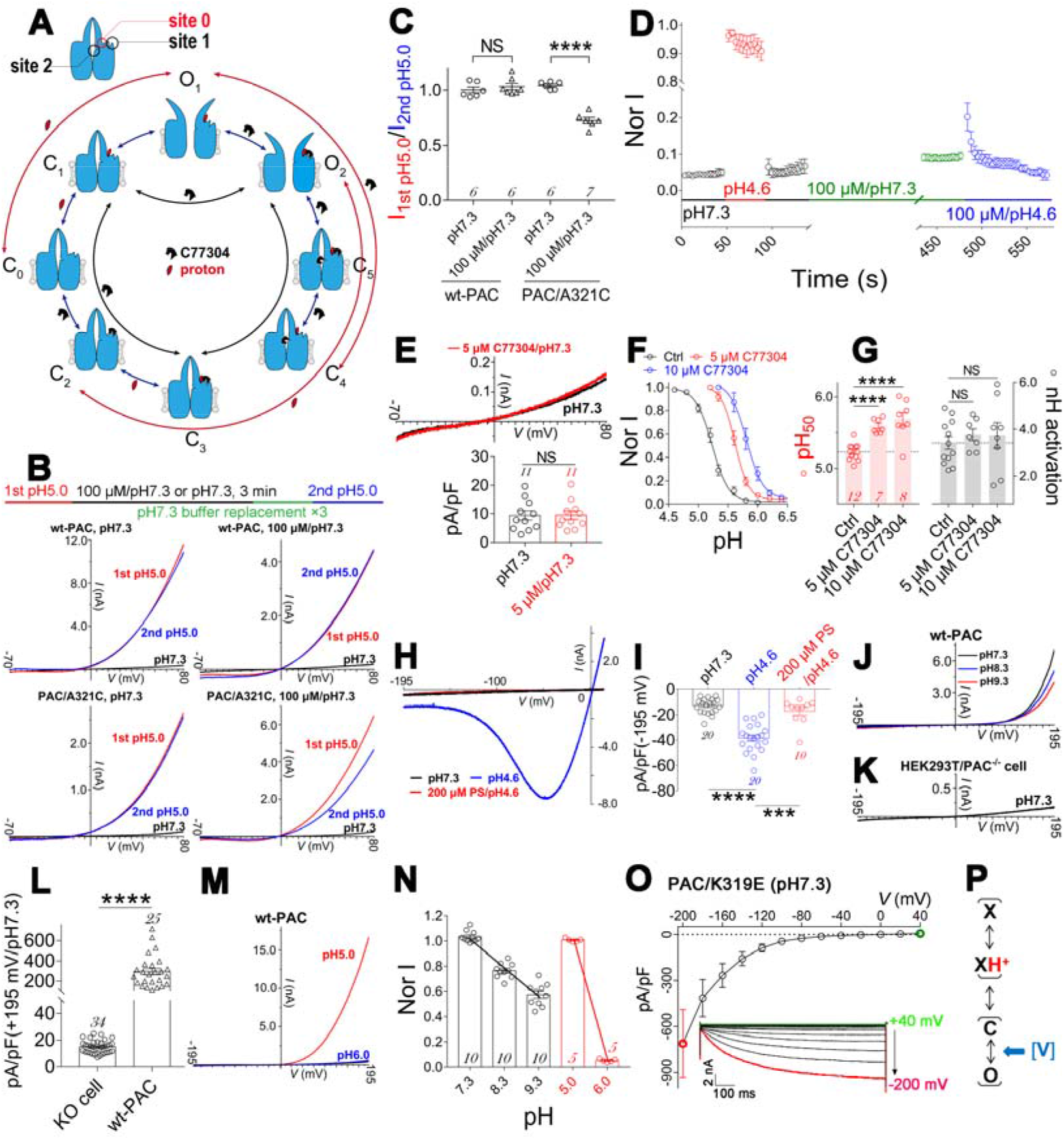
The mechanism of C77304 acting on the PAC channel. (A), Scheme illustrating the binding/unbinding of protons and/or C77304 with PAC channel and the resulting state transitions. ‘C_0_-C_5_’ and ‘O_1_-O_2_’ represent different closed and open states, respectively. site 0, proton binding; site 1, activation site; site 2, inhibition site. (B), Upper panel: experimental protocol; lower four panels: representative traces showing 100 μM C77304 pre-incubated at pH7.3 inhibits currents from A321C mutant, but not wt-PAC, channels (n = 6-7). (C), Summary of data in panel B (n value as indicated in each bar; NS, not significant; ****, *p*<0.0001; unpaired t-test). (D), Time course of preincubated C77304 (100 μM/pH7.3 for 5 min; see sequence of treatment conditions) blocking PAC currents upon channel activation by pH4.6 (the blue circles) (n = 7). (E), Example traces (upper panel) and summary analysis (lower panel) showing C77304 did not activate PAC currents at pH7.3 (n = 11; NS, not significant; paired t-test). (F)-(G), Current-pH relationships of PAC channels before and after 5 and 10 µM C77304 treatment (F) and the related summary analysis of the pH_50_s and slope factors (G) (n = 7-12; NS, not significant; ****, *p*<0.0001; ONE-WAY ANOVA with post-hoc Dunnett analysis). (H)-(I), Representative traces (H) and summary data (I) showing pH4.6 treatment activated a small but significant PS-sensitive and PAC channel-responsible currents at hyperpolarizing voltages, currents were elicited by ramp depolarization from -195 mV to +10 mV (n = 10-20; ****, p<0.0001; ***, p<0.001; paired t-test). (J)-(L), Representative current traces (J and K) and summary bar graphs (L) showing strong ramp depolarization (−195 mV to +195 mV) elicits large currents in wt-PAC transfected but not PAC knock-out cells at pH7.3, with extracellular alkalization to pH8.3 and 9.3 slightly reducing the amplitude (J) (n = 10-34; ****, p<0.0001; unpaired t-test). (M), Representative current traces showing pH6.0 bath perfusion dramatically reduced the pH5.0 acidification elicited currents in response to a -195 mV to +195 mV ramp depolarization (n = 5). Untransfected HEK293T cells expressing endogenous PAC channel were used for controlling the current amplitude. (N), Current-pH relationships of PAC channels in different pH ranges, with the linear fit slope factor determined to be -0.22 and -0.95 between pH7.3-9.3 and pH5.0-6.0, respectively; n value as indicated in each bar. Currents were recorded as in (J) and (M). (O), I-V relationship of PAC/K319E mutant channel at pH7.3, the currents were elicited by a cluster of voltage step from +40 mV to -200 mV (1 second) from the holding potential of 0 mV, inset shows a family of representative current traces (n = 5). (P), gating scheme of the PAC channel, C and O represent the closed and open pore, X and X(H^+^) represent the proton sensor with or without proton binding, respectively; and voltage (V) gates the pore through the conducting ions.

C77304 did not potentiate PAC currents at pH7.3 (Fig. 2E), suggesting a proton gating-dependent activation (i.e., C_2_→O_2_ transition is invalid without proton binding; Fig. 2A). Indeed, C77304 concentration-dependently shifted the channel’s current-pH relationship to the alkaline direction without changing the slope factor (Figs. 2F-G, Supplementary Fig. 3A, Supplementary Table 1), implying it increased the channel’s apparent proton affinity by facilitating proton binding (promoting C_2_→C_3_→O_2_ and C_2_→O_2_ transitions; Fig. 2A) or enhancing proton gating (promoting C_1_→O_2_ and C_1_→C_3_→O_2_ transitions; Fig. 2A), which were difficult to discriminate between. We eliminated the possibility of C77304 inhibiting the PAC channel by driving its desensitization as the PAC/A321C mutant was directly inhibited at pH7.3 but bidirectionally modulated at acidic pH6.2 (Supplementary Figs. 3B-C).

The outwardly-rectifying property of PAC currents means that depolarization increases the channel’s opening probability, which is mechanistically attributable to voltage-dependent protonation or protonation-independent voltage gating. Our data supports the latter, since: (i) at pH4.6, PAC channels were activated at very hyperpolarizing voltages and mediated small PS (pregnenolone sulfate^23^, a PAC channel inhibitor)-sensitive inward currents (Figs. 2H-I), suggesting the channels bind to protons voltage-independently and exhibit a small intrinsic opening probability when solely protonated; (ii) at pH7.3, PAC channels were activated by strong ramp depolarizations from -195 mV to +195 mV, conducting large outwardly-rectifying currents, which were absent in the PAC knock-out cells (Figs. 2J-L). Moreover, reducing pH from 7.3 to 8.3 and pH9.3, respectively, only slightly attenuated the currents (Figs. 2J), whereas reducing the proton concentration by 10-fold from pH5.0 to 6.0, within the pH range where PAC channels were typically gated by protons, nearly eliminated the currents (Fig. 2M), with the slope factor of the current-pH relationship by linear fit being determined as -0.22 and -0.95, respectively (Fig. 2N). We surmise that the current rundown caused by alkaline pHs (8.3, 9.3) results from non-specific damage to plasma membrane components. (iii) switching the conducting ion from Cl^-^ to Na^+^ by the K319E mutation^11^, converted the channel to be inwardly-rectified (Fig. 2O), suggesting the voltage dependence on the permeable ion species but not the channel protein itself, thereby supporting an ion-flux gating mechanism, as in the K2P channel^26^. Consequently, a refined gating scheme for the PAC channel is depicted in Fig. 2P, in which the voltage gating (V) and proton gating (X↔XH^+^) work, both separately and synergistically, to drive pore opening (C↔O). C77304 inhibited, but did not activate, the strong-depolarization activated PAC currents (Supplementary Fig.3D), supporting its role in potentiating channel activation via actions on proton-but not the voltage-gating.

These data implicate an intrinsic proton gating-independent voltage gating machinery in the PAC channel. The model in Fig. 2A fully recapitulate the bidirectional modulation of C77304 on the PAC channel, with C77304 binding to sites 1 and 2 with differing affinities and state-dependence to regulate channel gating. Furthermore, the potentiating effect of C77304 was more pronounced at pHs at the onset (‘root’) of the current-pH curve (hereafter, ‘root pHs’), wherein most channels were in the resting state.

### Mapping key residues involved in the proton gating of PAC channel and uncovering activation site 1

C77304 could be used as the pharmacological probe to identify the gating-related residues in PAC as it potentiated PAC currents only at root pHs. In this scenario, at pH5.0, C77304 at 5 µM should activate, but not inhibit, those mutants with acidic-direction shifted threshold activation pHs. Indeed, among the 28 alanine-scan mutants of the titratable residues in PAC channel, the H130A, D200A, E209A, E224A, E249A, E250A, D251A, E286A, and D289A mutants were activated in this experimental setting (Fig. 3A, Supplementary Fig. 4A), suggesting an attenuation of their apparent proton sensitivity. Their current-pH relationships were shifted to the acidic direction to different extents, with remarkably reduced pH_50_ but unchanged slope factor when compared to wt-PAC (Fig. 3B, Supplementary Figs. 4B, Supplementary Table 1), which reciprocally validated our hypothesis. Most importantly, we identified three glutamate residue mutations, E181A, E257A, and E261A, that fully abolished the proton gating of PAC (see Fig. 4).

**Fig 3.**
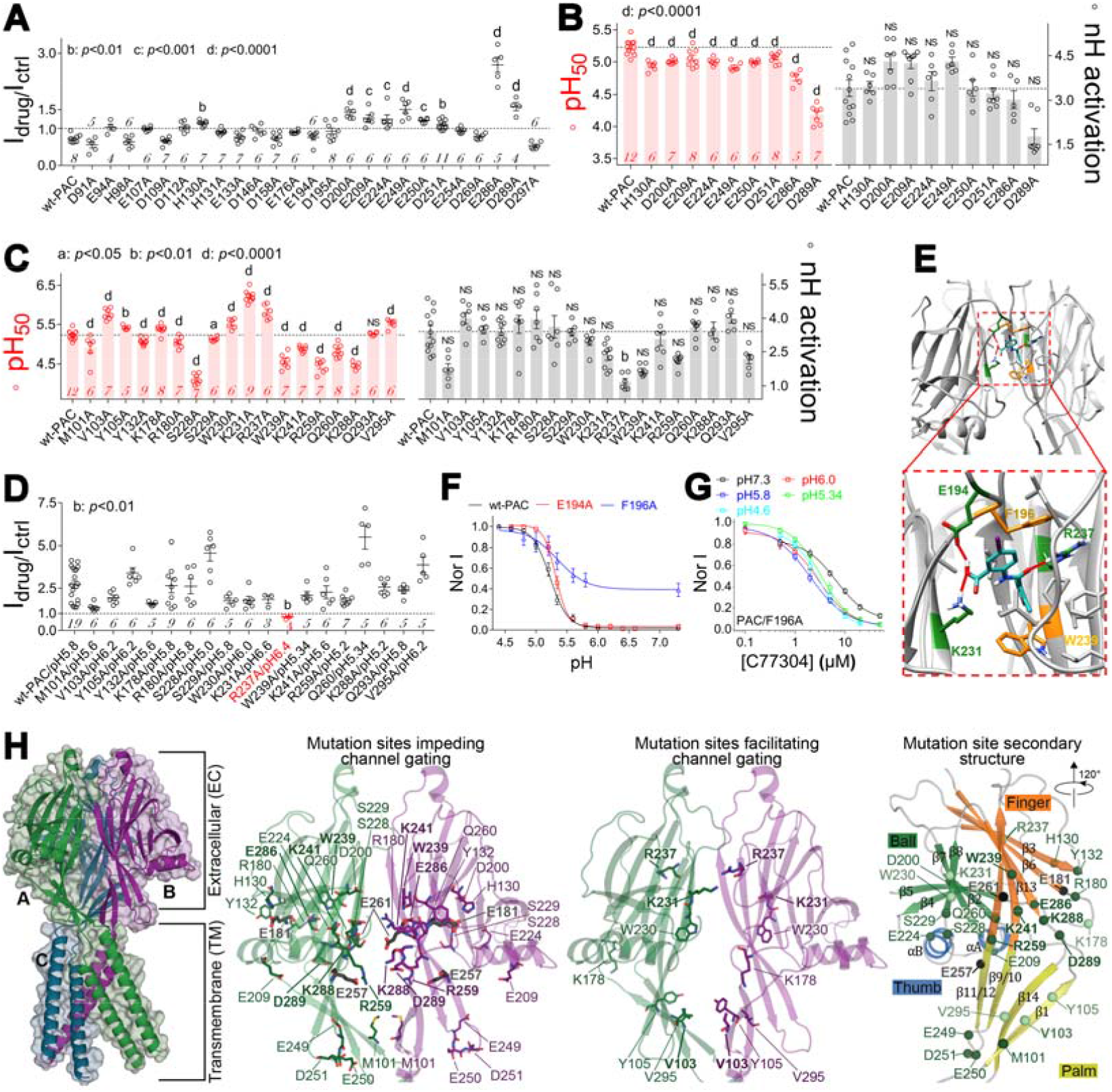
Mapping the key residues involved in proton gating and deciphering the activation site 1 in PAC channel. (A), Alanine-scanning mutations of glutamate, aspartate, and histidine residues in human PAC channel, bar graph shows the normalized effect of 5 µM C77304 on their currents at pH5.0. (B), Summary analysis showing the changes in pH_50_ and nH activation values of PAC mutants activated by C77304 in (A). (C), Summary analysis showing most mutations in the side portal region of PAC channel significantly change the pH_50_ of proton activation, and the R237A mutation remarkably changed the slope factor. (D), Effects of 5 µM C77304 on PAC channel mutant currents at their respective root pHs; currents were normalized to their respective control currents before drug treatment, showing the compound exclusively inhibited the PAC/R237A currents but activated the others. (E), Molecular docking of C77304 to PAC channel around the R237 site revealed that E194, F196, K231, R237, and W239 form the binding pocket. (F), Current-pH relationships of PAC/E194A and PAC/F196A mutant channels, wt-PAC was included for comparison (n = 6-12). (G), Dose-response relationships of C77304 inhibiting PAC/F196A mutant channel at multiple pHs spanning its current-pH relationship. The IC_50_s and mean slope factors were determined to be 6.6 ± 0.9 µM and 1.4 (n = 8), 2.9 ± 0.2 µM and 1.3 (n = 6), 2.0 ± 0.2 µM and 1.2 (n = 7), 3.6 ± 0.3 µM and 1.5 (n = 5), 3.0 ± 0.5 µM and 1.4 (n = 7), at pH7.3, pH6.0, pH5.8, pH5.34 and pH4.6, respectively. (H), PAC channel structure (PDB entry: 7SQG) showing locations of mutation sites as indicated. Bold text indicates mutations that had the strongest effects. At far right, domains and secondary structure elements are labeled according to PMID: 35878032. In (A) - (D), the differences between mutant and wt-PAC channels were assessed by ONE-WAY ANOVA with post-hoc Dunnett analysis. *p* values as indicated in each panel and ‘NS’ means not significant, n values for each channel as indicated in the bar.

**Fig 4.**
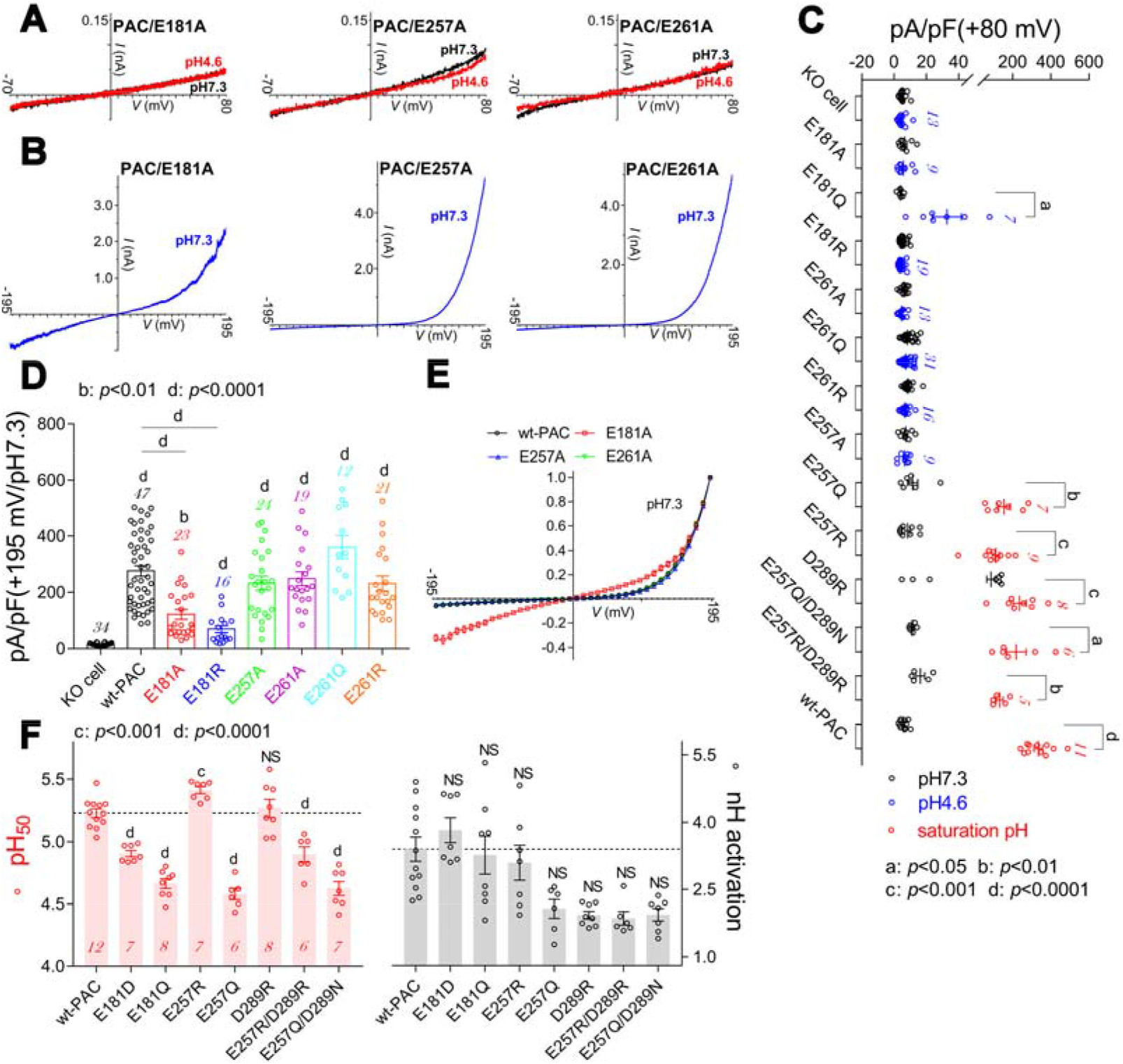
Characterizing the primary proton sensors in human PAC. (A)-(B), Representative current traces showing PAC/E181A, PAC/E257A, and PAC/E261A mutant channels did not respond to low pH_o_ stimulation (A) but were activated by strong ramp depolarizations from -195 mV to +195 mV at pH7.3 (B) (n = 9 - 24). (C)-(D), Summary analysis of proton (C) and strong depolarization (D; at pH7.3) activated currents of mutant channels as indicated, the KO cell and wt-PAC groups were included for comparison (n value as indicated in each bar). (E), Normalized I-V relationships of wt-PAC, PAC/E181, PAC/E257, and PAC/E261A mutant channels, with E181A mutation changed the I-V shape (n = 8-12). (F), Summary analysis of pH_50_ (left panel) and nH activation (right panels) values for mutant channels as indicated (n values inside each bar). Statistical differences were assessed using paired t-test (C); ONE-WAY ANOVA with post-hoc Dunnett analysis in (D) and (F); in (D), mutants were compared with KO and wt-PAC respectively. *p* values as indicated in each panel and ‘NS’ means not significant. Note in (D), the ‘NS’ symbols between E257A, E261A, E261Q, E261R and wt-PAC were omitted for clarity.

Molecular docking of C77304 to the PAC channel (PDB: 7JNA^11^) revealed its preferred residence in the channel’s side portal region. Consequently, we mutated residues comprising the side portal to alanines and measured their current-pH relationships, thus enabling determination of their respective root pHs for mapping the residues comprising activation site 1. Most mutations profoundly affected channel gating: M101A, Y132A, R180A, S228A, S229A, W239A, K241, R259A, Q260A, and K288A mutations significantly reduced the channel’s pH_50_; V103A, Y105A, K178A, W230A, K231A, R237A, and V295A mutations increased pH_50_ (Fig. 3C, Supplementary Fig. 5A, Supplementary Table 1). The R237A mutation also remarkably decreased the slope factor of the current-pH relationship (Fig. 3C).

Mutating key residues in site 1 is expected to eliminate the activation effect of C77304. In fact, currents through PAC/R237A mutant, but not any others, were significantly inhibited when tested at their respective root pHs (Fig. 3D, Supplementary Fig. 5B). Furthermore, this inhibitory effect was also observed at multiple pHs tested (Supplementary Fig. 5C). Subsequent docking of C77304 to the channel around R237 predicted four additional candidate binding residues - E194, F196, K231 and W239 (Fig. 3E). Among them, K231 and W239 mutations did not perturb the activating effect of C77304 (Fig. 3D), suggesting that these were not key for anchoring C77304 in the pocket. Interestingly, at their respective root pHs, the F196A, but not the E194A, mutation prevented activation by C77304, as seen with R237A (Fig. 3F, Supplementary Fig. 5C). The concentration-response relationship of C77304 against the F196A mutant further confirmed that it monotonically inhibits the currents at multiple pHs spanning the current-pH relationship curve (Fig. 3G). As expected, combining the R237A and F196A mutations also abolished C77304 activation (Supplementary Figs. 5D-E). In summary, these data show that F196 and R237 form activation site 1 in the PAC channel.

Mapping these gating-related residues to PAC structure revealed their condensed distribution in the middle ECD, surrounding E181, E257, and E261 (Fig. 3H). Most of these residues are not titratable at physiological pH and thus are unlikely the proton sensors. Instead, they might subtly adjust the biochemical environments where the proton sensors reside in to set their *p*K_a_, or affect the gating of protonated channels.

### Deciphering the primary proton sensors in PAC channel

The protonation of proton sensors in the PAC channel triggers its activation; therefore, mutating key proton sensing residues should abolish the channel’s proton gating by depleting its proton binding capability. Indeed, E181A, E257A, and E261A mutations eliminated PAC channel proton activation (Figs. 4A, 4C). Immunocytochemistry experiments revealed that these mutant channels were targeted to cell membranes much like wt-PAC (Supplementary Fig. 6A). Intriguingly, their voltage-dependent activation at pH7.3 was not impaired, with peak current density for both the E257A and E261A mutants being similar to wt-PAC (Figs. 4B, 4D). Moreover, the I-V relationships of E257A and E261A mutants mirrored that of wt-PAC, suggesting unchanged voltage sensitivity (Fig. 4E). The E181 mutation, however, significantly altered the I-V relationship and reduced voltage-activated currents when compared with wt-PAC, implying that it’s also involved in the channel’s trafficking and/or voltage-dependent gating (Figs. 4D-E).

Residue E257 was proposed to be involved in proton sensing in human PAC, via its interaction with D289 from the neighboring subunit^13^. Thus, we examined the potential role of the E257-D289 pairing in the channel’s proton gating using a multiple-attribute mutation analysis strategy (mutating E257 and D289 to R, Q or N). All these mutants were functionally gated by protons (Fig. 4C, Supplementary Fig. 6B). Consistent with a previous study^13^, the E257Q mutation strongly shifted the current-pH relationship to the acidic direction (Fig. 4F, Supplementary Fig. 6C, Supplementary Table 1). We reasoned that stabilizing the E257-D289 interaction would facilitate, while destroying it would impede channel gating. Indeed, the E257R mutation significantly shifted the current-pH relationship to the alkaline direction, whereas the D289R mutation produced a large basal opening at pH7.3 without a pH_50_ shift (Fig. 4F, Supplementary Fig. 6C, Supplementary Table 1). In contrast, the E257Q/D289N as well as the E257R/D289R double mutations caused a remarkable acidic-direction pH_50_ shift (Figs. 4F, Supplementary Fig. 6C, Supplementary Table 1). The E257R/D289R mutations were predicted to destroy the proposed pairing to eliminate channel gating; however, this was not observed experimentally. Therefore, these data argue that the E257-D289 interaction is not the primary driving force for PAC channel’s proton activation, but may facilitate channel gating.

Like E261A, the E261Q and E261R mutant channels were not functionally gated by protons (Fig. 4C, Supplementary Fig. 6D), and their voltage-dependent gating was not impaired (Fig. 4D, Supplementary Figs. 6E-F). The multiple-attribute mutations of E181 exhibited an intermediate phenotype compared to the E257 and E261 mutations: the E181Q mutant was normally gated by protons but with significantly reduced pH_50_ and current density (Figs. 4C, 4F, Supplementary Figs. 6G, 6I, Supplementary Table 1); the E181R mutant channel was refractory to proton gating with its voltage-dependent gating unaffected (Figs. 4C-D, Supplementary Fig. 6H). Unexpectedly, the E181D mutation also caused a remarkable acidic-direction pH_50_ shift (Fig. 4F, Supplementary Fig. 6I, Supplementary Table 1), suggesting the side chain length might affect the channel’s proton gating as well. We conclude that E181, E257 and E261 contribute to the channels’ primary proton sensors, although their individual role in channel gating might vary.

### Conservation and variation of the proton sensing mechanisms among orthologous PAC channels

The PAC channel is evolutionarily conserved across the animal kingdom. To further validate the importance of the identified proton sensors in human PAC, we performed cross-species analysis by mutating their analogous sites in various orthologous PAC channels from naked mole rat *H. glaber* (hgl-PAC), eastern brown snake *P. textilis* (pte-PAC), the Indian cobra *N. naja* (nna-PAC), red junglefowl *G. gallus* (gga-PAC), *and* zebrafish *D. rerio* (dre-PAC). As expected, they all functionally conducted proton-evoked (left panels in Figs. 5A-E, Supplementary Figs. 7A, 7D, 7G, 7J and 7M) as well as voltage-activated (at pH7.3) currents (right panels in Figs. 5A-E, Supplementary Figs. 7A, 7D, 7G, 7J and 7M). Excitingly, mutating the E257 and E261 analogous sites in hgl-PAC, pte-PAC, and nna-PAC channels abolished their proton gating without affecting the voltage-dependent gating (Figs. 5A-C, Supplementary Figs. 7B-C, 7E-F and 7H-I), which confirms the critical role of these two residues in proton sensing of PAC channels across species. The E181 analogous site mutation in pte-PAC and nna-PAC channels specifically eliminated proton gating as well (Figs. 5B-C, Supplementary Figs. 7E-F and 7H-I). Unexpectedly, the E181 analogous site mutation in hgl-PAC did not fundamentally affect its proton gating, although reduced the current density (left panel in Fig. 5A, Supplementary Fig. 7B). Furthermore, mutating E181, but not E257 and E261, residues in gga-PAC (E182A) and dre-PAC (E183A) specifically abolished proton gating (Figs. 5D-E, Supplementary Figs. 7K-L and 7N-O). These data highlight conservation and variation in the proton sensing machinery of orthologous PAC channels (Fig. 5F). An extensive scanning-mutation analysis of all titratable residues in dre-PAC also proved that the E183A mutation exclusively eliminated the channel’s proton gating (Fig. 5G). Therefore, the variation of the residues constructing the proton sensors in orthologous PAC channels is likely limited to three key residues– E181, E257 and E261 – identified in human PAC (Fig. 5F), with one or two key sites being abandoned during evolution in a species-specific manner.

**Fig 5.**
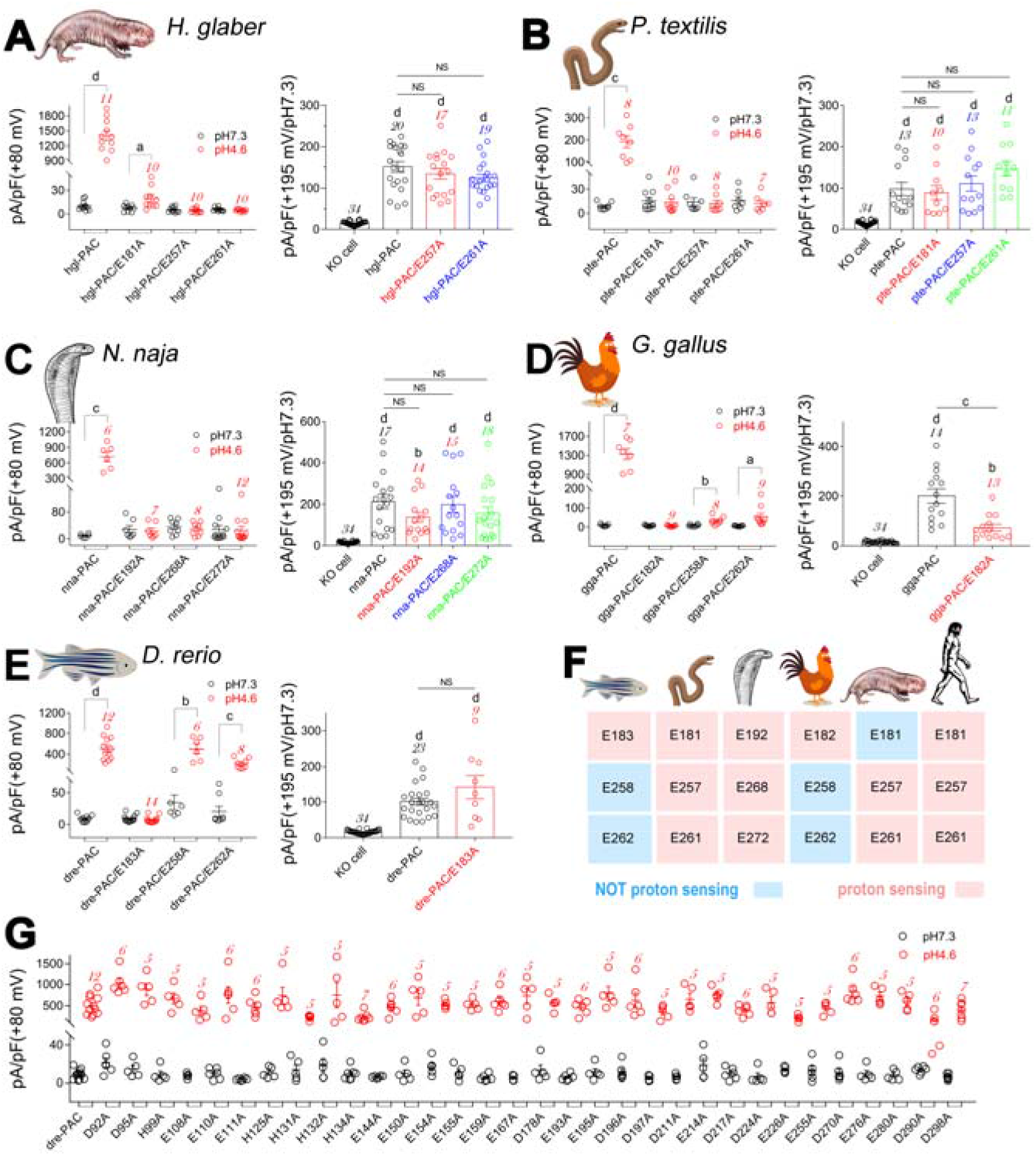
Cross-species analysis of the proton sensing mechanism of PAC channel. (A), Summary analysis of the proton (left) and strong-depolarization (right) activated currents of hgl-PAC, hgl-PAC/E181A, hgl-PAC/E257A, and hgl-PAC/E261A mutant channels from *H. glaber*. (B), Summary analysis of proton (left) and strong-depolarization (right) activated currents in pte-PAC, pte-PAC/E181A, pte-PAC/E257A, and pte-PAC/E261A mutant channels from *P. textilis*. (C), Summary analysis of proton (left) and strong-depolarization (right) activated currents of nna-PAC, nna-PAC/E192A, nna-PAC/E268A, and nna-PAC/E272A mutant channels from *N. naja*. (D), Summary analysis of proton (left) and strong-depolarization (right) activated currents in gga-PAC, gga-PAC/E182A, gga-PAC/E258A, and gga-PAC/E262A mutant channels from *G. gallus*. (E), Summary analysis of proton (left) and strong-depolarization (right) activated currents in dre-PAC, dre-PAC/E183A, dre-PAC/E258A, and dre-PAC/E262A mutant channels from *D. rerio*. (F), Heat map showing the homology of residues constituting the presumptive proton sensors among orthologous PAC channels. (G), Alanine scan mutation of titratable residues in dre-PAC channel; all of these mutants were effectively activated by protons by pH dropping from 7.3 to 4.6. Currents were recorded with -70 mV to +80 mV ramp depolarizations. Statistical differences between pH7.3 and pH4.6 in (A) - (E) were assessed by paired t-tests; differences for strong-depolarization activated currents between groups in (A) - (E) were assessed by ONE-WAY ANOVA with post-hoc Dunnett analysis (the KO cell and wt-PAC groups were used as control for comparison, respectively); a, p<0.05; b, p<0.01; c, p<0.001; d, p<0.0001; NS, not significant. n value as indicated in each bar.

### A gating model of PAC channel as revealed by MD simulations

MD simulations show that the ECD of PAC channel in the resting state contains a complex network of interactions within subunits (Figs. 6A-B). For example, E261 interacts stably with K288 from the same subunit (Fig. 6C). In the open state, the electrostatic repulsion between subunits is decreased, partially due to protonation of E261 (and probably others) at pH4, leading to a significant decrease in intra-subunit interactions and increase in inter-subunit interactions. Consequently, a less expanded ECD was observed, consistent with its cryo-electron microscopic structure^11^.

**Fig 6.**
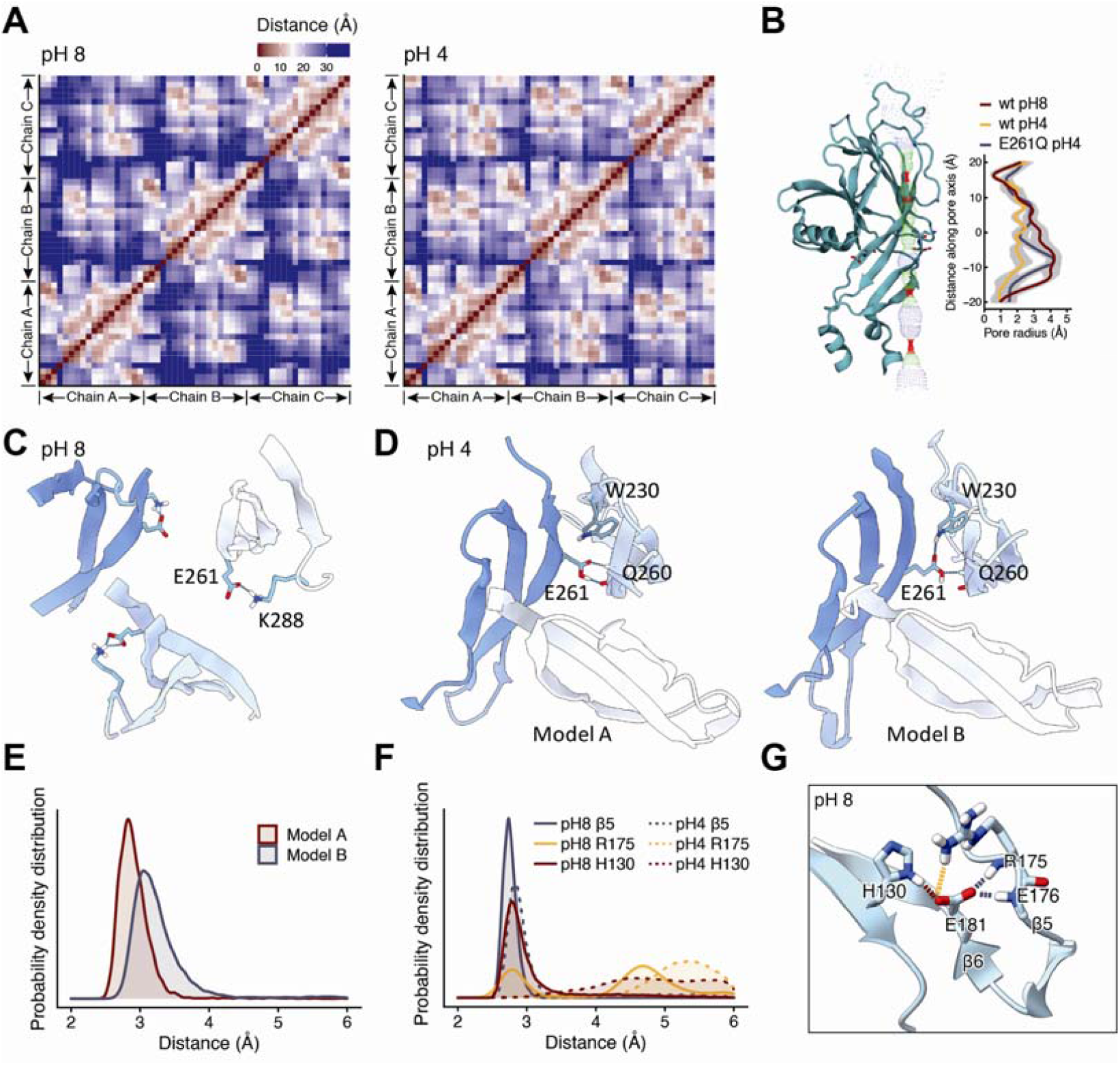
Building a gating model of the PAC channel. (A) Inter- and intra-chain contact maps at pH = 8 (left panel) and pH = 4 (right panel). The x/y axis represents the residue ID on the three chains. The color bar indicates the distance (in angstroms) of the interactions. (B) The pore sizes of the wild-type PAC channel in the resting and activated states, and the E261Q mutant PAC channel in the activated state. Only one of three subunits is shown for clarity. (C) The E261 interaction pattern in a representative resting state structure, dominated by electrostatic attractions within the subunit. The colors indicate the three subunits. (D) The interactions between Q260 and the E261-W230 cluster on a neighboring chain in the activated state. The different orientations of E261 lead to the existence of models A (*left*) and B (*right*). (E) The probability density distributions of the distance between the carboxyl oxygens on E261 and on Q260 in the two models. (F) The probability density distributions of the hydrogen bond formed with E181 under different conditions. (G) The interaction network involving E181 in a representative structure of a wild-type PAC channel in the resting state taken from the MD trajectory. The color representation of different hydrogen bonds is consistent with that used in panel F.

The neutralized E261 hydrogen bonds with Q260 from the adjacent subunit (Model A, Fig. 6D). Notably, the E261-Q260 interactions are located at the neck of the ECD cavity, where its contraction begins in the active state (Fig. 6B) and is essential for conformational transition. Simulations show that E261 can also interact with W230 in the neighboring subunit (Model B in Fig. 6D), thus interfering with the E261-Q260 link by pushing E261 outward (Fig. 6E), rendering difficult the conformational transition of ECD upon protonation. In contrast, the W230A mutation eliminates this interference; these data are congruent with the W230A mutant being more sensitive to acidic environments (Fig 3C).

We further found that this interaction cannot be maintained in the E261Q mutant, and therefore there was no ECD contraction (Fig. 6B). This observation is consistent with experiment showing that the E261A/E261Q mutants lack acidic responses (Fig. 4C). Presumably, the conformational rearrangement resulted from the altered protonation state of E261 is an integral part of its acid-sensitive mechanism, whereas the non-titrating residues are unable to play such a role.

E181 in the β6 strand of PAC channel was another crucial residue. MD simulations show that the side-chain of E181 stably forms a two-dentate interaction with the backbone of E176 and R175 on β5 (Fig. 6F-G). These interactions are essential for maintaining the coupling between β5 and β6 and are important links in the transmission of conformational changes. The E181D mutant with a shorter side-chain should weaken this coupling, making it less sensitive to acid. In addition, the charge-reversal E181R mutant was supposed to result in a complete loss of interaction. Both scenarios were proven correct in our experiments (Figs. 4C and 4F).

## Discussion

C77304 is a novel and unique bidirectional modulator of the PAC channel. That a modulator bi-directionally regulates ion channels is not unprecedented, as observed in the modulations of NaV1.3 by PF-06526290 and NaChBac by the general anesthetic drug sevoflurane^27,28^. One common feature of these promiscuous compounds is that they bind to different interacting sites state-dependently. Although C77304 binds exclusively to site 2 in PAC channel’s open state, it may activate the channel by binding to either the closed state to promote proton binding (C_2_→C_3_→O_2_ and C_2_→O_2_ transitions; Fig. 2A), the proton-bound pre-open state to enhance the pore opening (C_1_→O_2_ and C_1_→C_3_→O_2_ transitions; Fig. 2A), or even the open state to stabilize channel opening (O_1_→O_2_ transition; Fig. 2A). We were unable to discriminate the binding effect from the gating effect^29^, both of which could contribute to the facilitated proton gating by C77304. Accordingly, we incorporated all these mechanisms to build a gating model (Fig. 2A). Importantly, C77304 can detect pH_50_ shifts as small as ∼0.15 pH units (Figs. 3A-B), and is thus a valuable pharmacological probe for further dissecting proton gating in PAC.

Polymodal gating is commonly observed in ion channels: TRPV1, TRPM8, TRPA1 channels in the transient receptor potential channel superfamily, the calcium activated chloride channel TMEM16A, and the large-conductance calcium activated potassium channel (BK), are gated by more than one stimuli such as chemicals, acid, Ca^2+^, voltage, temperature, and mechanical force^30-34^. Membrane depolarization promotes PAC channel opening in acidic conditions resulting in outwardly-rectifying currents; however, whether membrane depolarization directly gates PAC was unknown. Here, we show that the PAC channel is also activated by strong depolarizations even at alkaline pHs, which positions it as polymodally gated.

As for the voltage-dependent gating of PAC, four theoretically possible scenarios exist: 1) voltage and proton individually and allosterically activate PAC channel, resembling the voltage- and Ca^2+^- dual-allosteric gating of the BK channel^35-37^; 2) allosteric modulation that utilizes the proton gating pathway, as in the TRPV1 channel, where voltage gating involves the proton gating machinery^38^, accordingly, protons cannot activate the channel at very negative hyperpolarizations; 3) voltage activates PAC channel by potentiating its proton binding, like in the TMEM16A channel^39^; and 4) voltage gates the channel through the conducting ions, that is the ion-flux gating mechanism as in the K2P channel. Structural analysis revealed no charged residues in the transmembrane electrical field outside the channel pore, which eliminated the possibility of pore control by a remote voltage sensor and thus scenario 1. The experimental data showed that protons effectively activated PAC channels even at very negative hyperpolarizations (Figs. 2H-I), which rules out scenario 2. Scenario 3 is invalidated as the pH-current relationships of PAC channels in acidic and neutral to alkaline pH ranges showed dramatically different slope factors (Fig. 2N), suggesting distinct gating mechanisms. Most importantly, converting PAC into a Na^+^-selective channel by the K319E mutation fully reversed its original outward rectification and rendered the channel inwardly-rectified (Fig. 2O), suggesting a voltage dependence on the conducting ion but not the channel protein itself, thereby strongly supporting scenario 4. Although the threshold activation voltage of ∼+100 mV at neutral pHs seems too high for activating PAC channels in physiological conditions, protons dramatically shifted the threshold voltage to near 0 mV, which adds importance to such voltage gating and suggests a crosstalk between voltage and proton gating. Moreover, it’s unknown whether heat can also facilitate voltage-dependent gating of PAC as a previous study showed that increasing temperature enhances the channel’s proton gating^40^. The coupling between voltage and proton gating, as well as the detailed mechanisms of voltage sensing, need further study. It’s likely that the residue constituting the selectivity filter in PAC, K319, acted as the voltage sensor, as in the K2P channel^26^.

The proton sensors in PAC remain undefined. As aforementioned, the Long group proposed that E257-D289, E249-E107, and E250-D297 pairing in acidic conditions confer proton sensing in human PAC^13^. In another study, the Qiu group proposed that H98, H130, H131, and D269 residues form the proton sensor, for which the H98R/H130R/H131R/D269A quadruple mutations greatly shifted the channel’s pH-current relationships to the alkaline direction accompanied by change of the slope factor and basal opening at neutral pH^41^. However, alanine-scanning mutagenesis of these residues (except for E257) did not eliminate the channel’s proton gating, which raises questions about their roles in proton sensing. Instead, we found that the E181A, E257A, and E261A mutations eliminated the channel’s proton gating (Fig. 4), arguing for their critical roles in proton sensing. The loss of proton-activated currents could also be attributed to the disruption of channel gating rather than proton sensing without evidence showing the mutants could be functionally gated, for example, by other stimuli. The Qiu group assigned E257 and E261 as components of a ‘joint region’ which acts to relay allosteric signals from the proton sensors^41^. Here, by revealing the voltage-dependent gating of PAC channel at neutral pHs, we showed that E181A, E257, and E261 mutations specifically eliminated the proton gating of PAC channel without affecting the voltage gating, therefore supporting the assertion that these residues are the primary proton sensors. Indeed, incorporating the E257A and E261A mutations into the PAC/A321C mutant also abolished the channel’s proton gating without affecting the proton-independent basal opening (Supplementary Fig. 8), which is further evidence of their critical roles in proton sensing. Moreover, disrupting the proposed E257-K288 interaction^41^ by the E257R mutation did not fundamentally affect the channel’s proton gating (Fig. 4C), therefore raising doubt as to its role in relaying gating signals. Cross-species analysis also validated the proton sensing machinery we identified in human PAC. Finally, sequence alignments showed that the E181, E257, and E261, but not the other gating-related residues are conserved across functional PAC channels from 13 different species (Supplementary Fig. 9). Collectively, the identified proton sensors and activation site 1 should be invaluable in inspiring future drug development targeting the PAC channel.

## Methods

### Compound synthesis

C77304 (5-iodo-2-(2-methylfuran-3-carboxamido)benzoicacid) was one ‘hit’ compound of the manual patch-clamp screening of PAC channel modulator from a chemical compound library (from Selleck Chemicals LLC, Catalog No. L3600), and was synthesized by Nafu Biotechnology (Shanghai Nafu Biotechnology Co., Ltd, Shanghai, China). The purity and structure of the synthetic C77304 was validated by HPLC and NMR analysis (Supplementary Figs. 1-2).

### Cell culture, plasmids and transient transfection

The HEK293T/PAC^-/-^ cell line was made by deleting a 31 bp long sequence(5’-AGCAGGACAAGGAGACGGTCAGAGTCCAAGG-3’) in exon 2 of the PAC coding gene PACC1 using the CRISPR-Cas9 method (Ubigene Biosciences, Guangzhou, China), the correct double knock-out the PACC1 gene was confirmed by PCR analysis and Sanger sequencing. KO cells were cultured in Dulbecco’s Modified Eagle’s Medium (DMEM) (Invitrogen; Thermo Fisher Scientific, Inc., Waltham, MA, USA) supplemented with 10% FBS and 1% PS (all from Gibco; Thermo Fisher Scientific, Waltham, MA, USA), maintained at 37 L in an incubator with saturated humidity and 5% CO_2_. PAC coding genes from different species (*Homo sampiens, Heterocephalus glaber, Pseudonaja textilis, Naja naja, Danio rerio, Gallus gallus, Anolis carolinensis, Latimeria chalumnae, Clupea harengus, Poecilia reticulata, Oreochromis niloticus, Nothobranchius furzeri*, and *Boleophthalmus pectinirostris*) were synthesized by Genscript (Genscript Corp., Nanjing, China) and cloned into the mammalian expression vector pCMV-blank. Channel mutants were made by site-directed mutations as previously described by us^42^. All constructs were sequenced to confirm correct mutations were made. Plasmids for wild-type or mutant PAC channels was transiently transfected into HEK293T/PAC^-/-^ cells using Lipofectamine per the manufacturer’s instructions (Invitrogen; Thermo Fisher Scientific, Inc., Waltham, MA, USA). The transfection amount for each construct was adjusted based on its functional expression level in cells. Six hours after transfection, cells were seeded onto poly L-lysine (PLL)-coated coverslips, and patch-clamp analysis was conducted 24-36 hours post transfection.

### Electrophysiology

Whole-cell patch clamp recordings were performed in an EPC-10 USB platform (HEKA Elektronik, Lambrecht, Germany). Recording pipettes with an access resistance of 2-3 MΩ after pipette solution filling were prepared from glass capillaries in a PC-10 puller (NARISHIGE, Tokoyo, Japan) using a two-steps program. To minimize pipette capacitance, only the tip of the pipette was filled with pipette solution. Artificial capacitance effect was reduced by sequential fast and slow capacitance compensation using the computer-controlled circuit of the amplifier. To minimize the leak current contamination, only cells with seal resistance higher than a gigaohm (GΩ) resistance after break in were selected for further analysis. Series resistance was kept to less than 10 MΩ to minimize voltage error, and 80% series resistance compensation was used with a speed value of 10 µs. Cells were held at 0 mV, and PAC currents were elicited by a voltage ramp from -70 mV to +80 mV with a speed value of 1 mV/ms (or -195 mV to +195 mV with a speed value of 0.5 mV/ms for recording the strong depolarization activated currents at pH7.3). Cell capacitance was measured using the LockIn Extention method when analyzing the current density. Recording solutions were as previously reported^8^: the extracellular solution contains (in mM): 145 NaCl, 2 KCl, 2 MgCl_2_, 1.5 CaCl_2_, 10 HEPES, 10 glucose (300 mOsm/kg; pH7.3 with NaOH); Different acidic pH solutions were made of the same ionic composition without HEPES but with 5 mM Na_3_-citrate as buffer and the pH was adjusted using citric acid; The standard pipette solution contains (in mM): 135 CsCl, 1 MgCl_2_, 2 CaCl_2_, 10 HEPES, 5 EGTA, 4 MgATP (280-290 mOsm/kg; pH7.2 with CsOH). All chemicals were purchased from Sigma-Aldrich (Sigma-Aldrich, Saint Louis, MO, USA). Drugs and solutions were applied by gravity perfusion. The concentration-response curves were fitted by a Hill logistic equation to estimate the potency (IC_50_ or EC_50_) of C77304. The pH-current relationships of wt-PAC and PAC mutants were fitted by the equation: Y=Y_(0)_ + (Y_steady_-Y_(0)_)/(1+10^((pH_50_-X)*nH)), in which pH_50_ and nH represents the half-activation pH and the slope factor of the curve, respectively.

### Immunocytochemistry

HEK293T/PAC^-/-^ cells transfected with human wt-PAC, PAC/E181A, PAC/E257A or PAC/E261A mutant channel (with enhanced green fluorescense protein (EGFP) fused to their C-termini) were seeded on PLL-coated coverslips, stained with DiI and DAPI, and fixed with 4% PFA (all from Sigma-Aldrich). The coverslips were mounted in nail polish and observed under IX83 inverted fluorescence microscope (Olympus corporation, TOKYO, Japan), fluorescence of cell membrane (red), channel protein (green) and the nucleus (blue) was sequentially elicited and merged using the cellSens Dimension software (version 2.3; Olympus corporation, TOKYO, Japan).

### System setup and molecular dynamics simulations

For resting state, the cryo-EM structure of PAC resting state (PDB entry: 7JNA^11^) was used. For open state, two cryo-EM structures (PDB entry: 7JNC^11^ & 7SQF^13^) were used. Since the experimental studies found that the mutation sites involved in the proton gating mechanism were concentrated in ECD, the ECD was extracted from the crystal structure to accelerate computational processing. The residues’ protonation states for the resting state in pH8.0 and the open state in pH4.0 were determined by the reported predicted pKa^11^. Next, the protein was embedded in water box. In addition to a few ions added to neutralize the system, a 150 mM concentration NaCl was added to mimic the physiological condition of salt concentration. This system contains a total of ∼11400 atoms, including ∼3200 water molecules.

For both systems, energy minimization was first used to remove bad contacts, where harmonic restraints were applied on the protein backbone atoms. The simulation system was then equilibrated, with gradually decreased harmonic position restraints applied to the heavy atoms of the protein backbone. After equilibration, the simulations were continued in the NPT ensemble at 1 atm pressure and 303 K for 400 ns. The first 50 ns of each trajectory was not used for analysis. We carried out two replicate MD simulations for each system and obtained consistent results. The MD simulations were carried out using the CUDA-accelerated NAMD program version 2.14^43^. The Charmm36 force field parameters^44^ and the TIP3P water model^45^ were used. Periodic boundary conditions were applied and the particle mesh Ewald method was used to treat long-range electrostatic interactions^46^. HOLE^47^ was used to calculated the distribution of pore radius. VMD^48^ and UCSF ChimeraX^49^ were used to analyze MD trajectories and visualize the models.

### Data analysis

Data were presented as MEAN ± SEM, n value was presented as the number of separate experimental cells. Data were analyzed using the softwares Igor (WaveMetrics Inc., OR, USA), Excel 2010 (Microsoft Corporation, Redmond, WA, USA), OriginPro 8 (Northampton, MA, USA) and Graphpad Prism (GraphPad Software, La Jolla, CA, USA). Statistics were analyzed using paired t-test, unpaired t-test or ONE-WAY ANOVA, comparisons between groups were performed using post-hoc analysis with the Dunnett method. Statistical difference was accepted at *p* < 0.05.

## Acknowledgements

This work was supported by the National Natural Science Foundation of China (Grant Nos. 31600669, 32071262, 31770832, 32171271, and 22273034), the Science and Technology Innovation Program of Hunan Province (2020RC4023), the Natural Science Foundation of Hunan Province (Grant No. 2018JJ3339), the Research Foundation of the Education Department of Hunan Province (Grant No.18B015), and the Fundamental Research Funds for the Central Universities. Parts of the calculations were performed using computational resources on an IBM Blade cluster system from the High-performance Computing Center (HPCC) of Nanjing University.

## Author contributions

C.T. designed experiments. P. Z., C. T., Y. Y., Z. X., G. L., H. L., Y. L., and Q. L. performed experiments. Y. Y. and H. D. performed molecular dynamics simulations. S. P.-M and H. Z. performed molecular docking. C. T., P. Z., H. D., and R. K. analyzed and interpreted data. F. Y. helped with interpreting the data. C. T., H. D., R. K., and Z. L. supervised the study and wrote the manuscript.

