## Supplementary Information for "A new polymodal gating model of the proton-activated chloride channel"

^6^Lead Contact

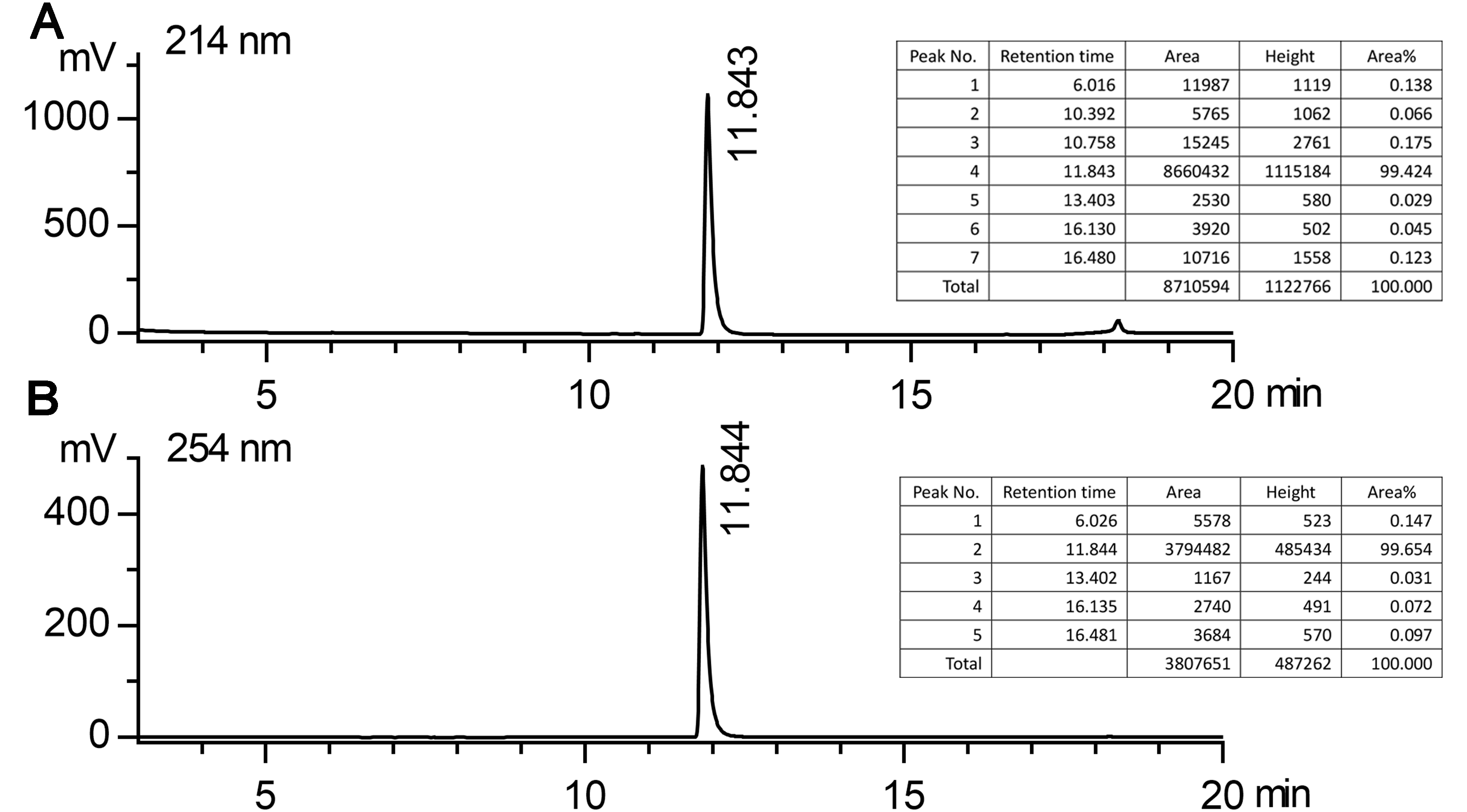

**Supplementary Fig. 1.** (A)-(B), Liquid-chromatography (LC) analysis demonstrated that the purity of synthesized C77304 is above 99% as determined by the peak area normalization method. (A), 214 nm; (B), 254 nm.

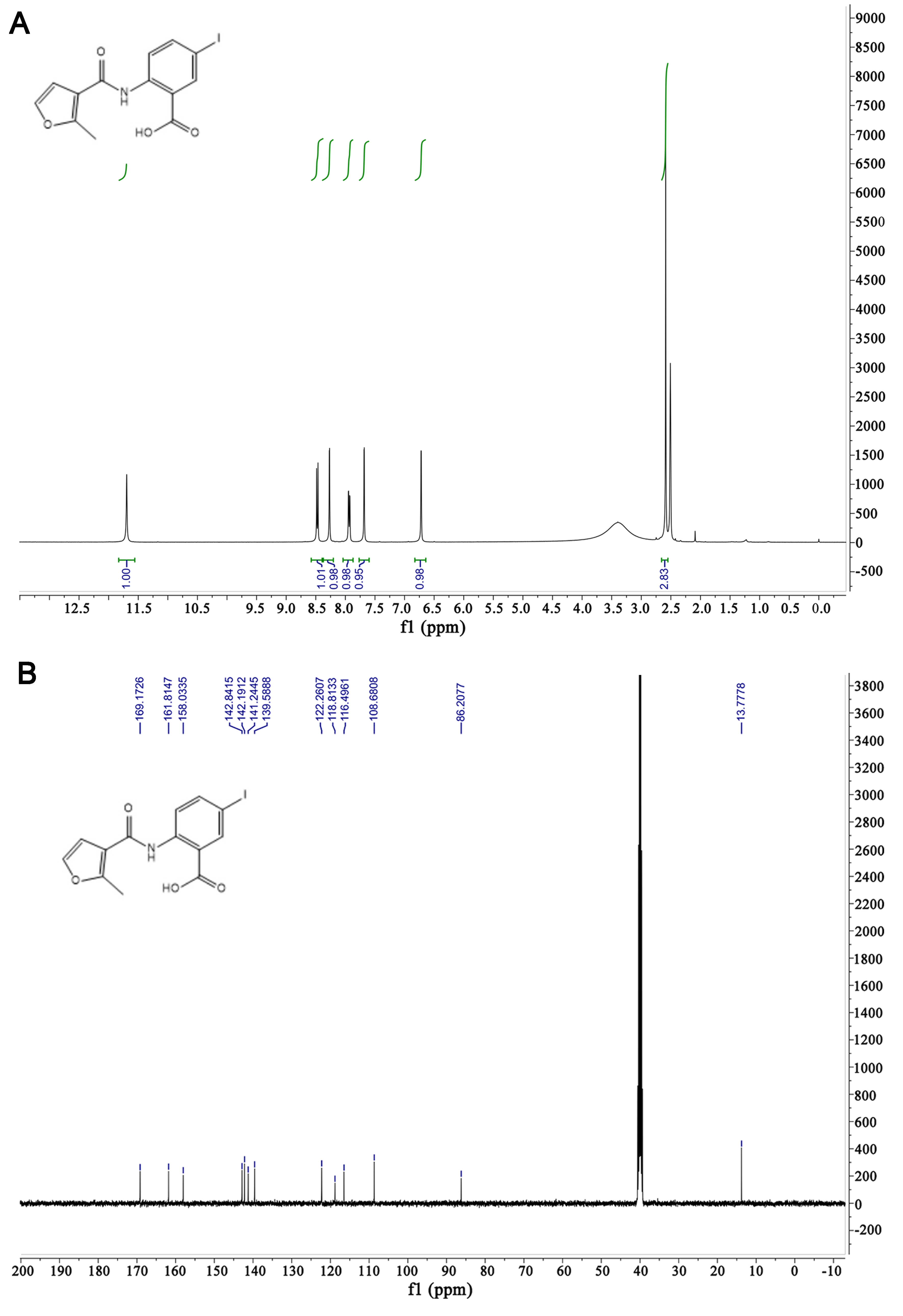

**Supplementary Fig. 2.** Structure validation of synthesized C77304 by ^1^H-NMR (A) and ^13^C-NMR (B).

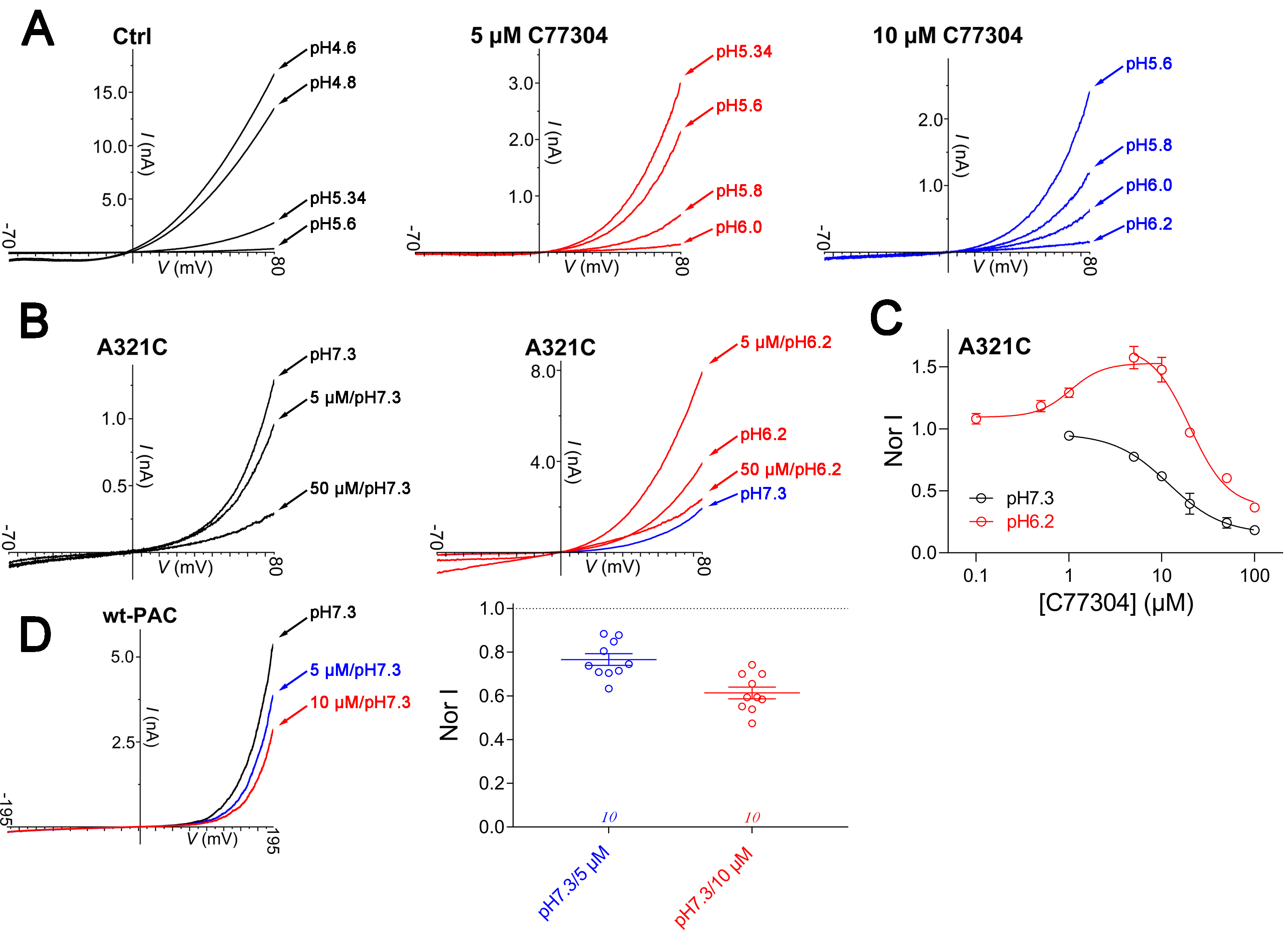

**Supplementary Fig. 3.** (A), Representative PAC currents elicited by different acidic pH solutions in the absence (*left*) and presence of 5 µM (*middle*) or 10 µM C77304 (*right*); note that C77304 treatment changed the threshold and saturating activation pHs. Currents were recorded with ramp depolarizations from -70 mV to +80 mV (n = 7 – 12 cells per condition). (B), Representative current traces showing C77304 monotonically inhibiting (*left*) or bidirectionally modulating (activating and inhibiting; *right*) PAC/A321C mutant channel at pH7.3 and pH6.2, respectively (n = 5). (C), Concentration-response relationships of C77304 acting on PAC/A321C mutant channel, with the curves being sigmoidally- or bell-shaped at pH7.3 and pH6.2, respectively. The EC_50_ for C77304 activating PAC currents at pH6.2 was 1.0 ± 0.4 µM, and the IC_50_ for inhibition were 11.8 ± 1.9 µM and 20.0 ± 2.9 µM at pH7.3 and pH6.2, respectively (n = 5). (D), Representative traces (left) and summary (right) showing C77304 dose-dependently inhibited strong depolarization activated PAC currents at pH7.3 (n = 10).

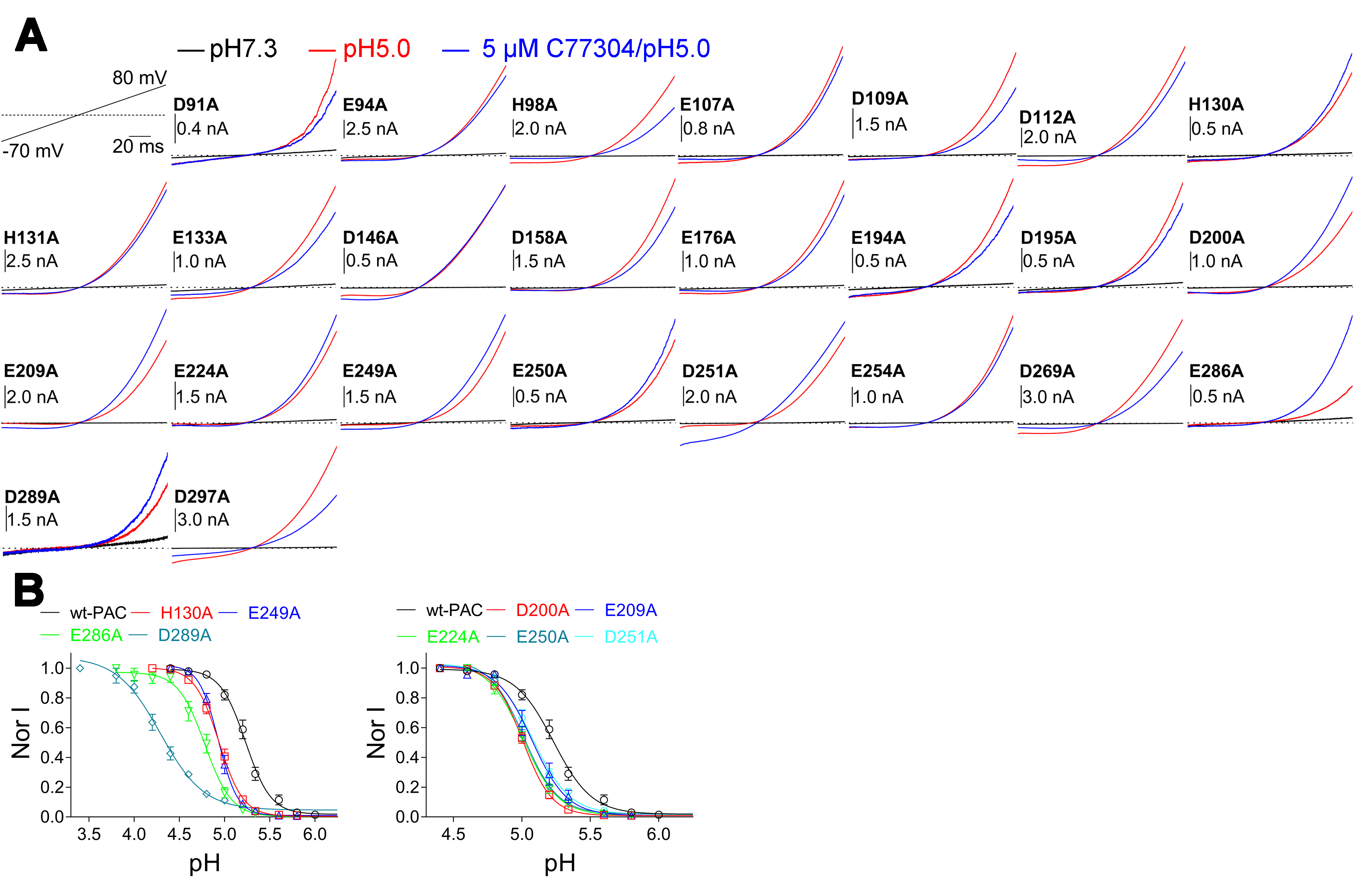

**Supplementary Fig. 4.** (A), Representative current traces showing the effect of 5 µM C77304 on PAC channel mutants of the titratable glutamate, aspartate and histidine residues at pH5.0; voltage protocol as shown (n = 4 – 11). (B), Current-pH relationships of the mutant PAC channels as indicated, showing differing extent of acidic direction shift compared with the wild-type (wt)-PAC (n = 5 - 12).

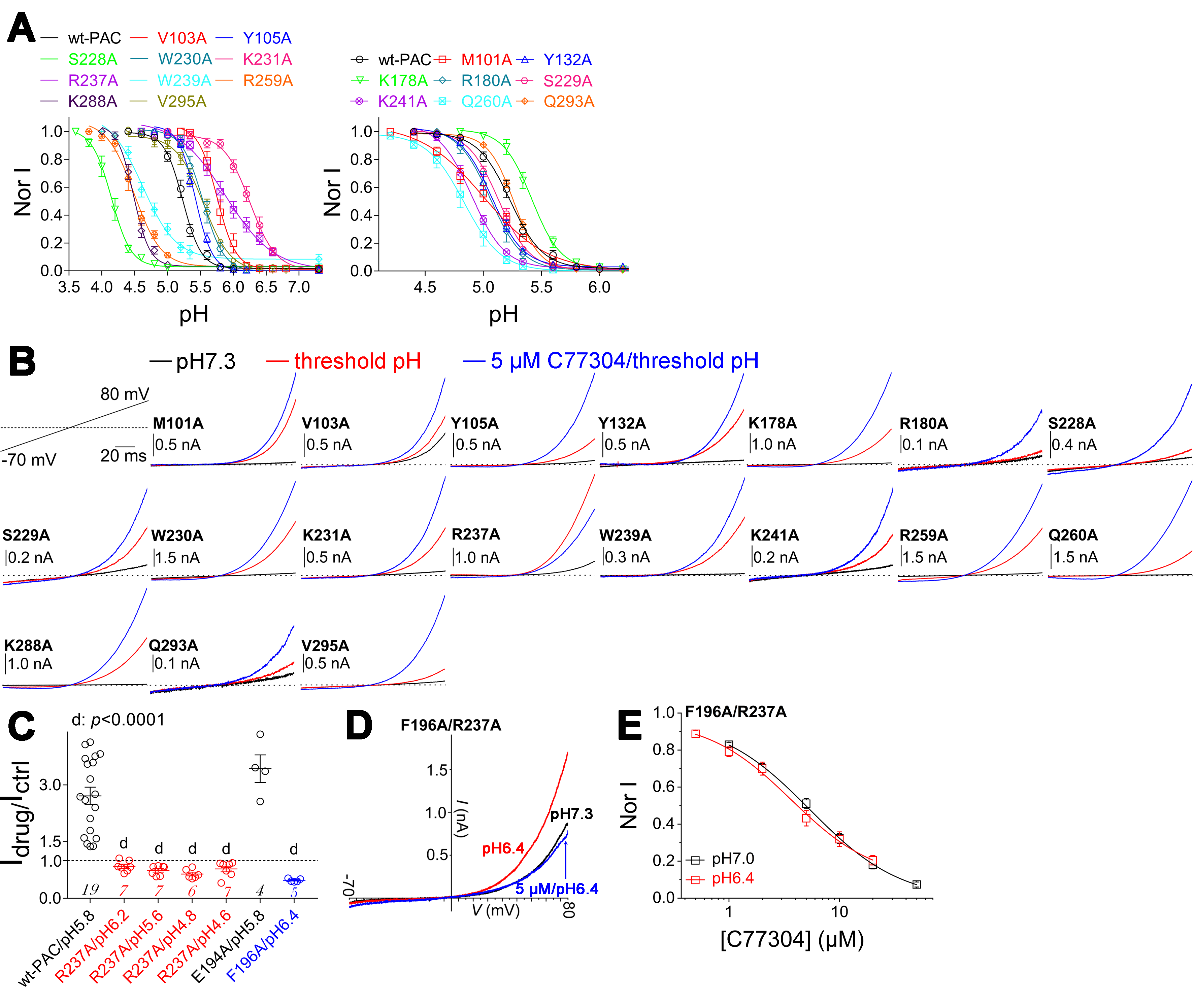

**Supplementary Fig. 5.** (A), Current-pH relationships showing that mutating residues in the side portal region of PAC channel affects its proton gating (n = 5 – 12). (B), Typical current traces demonstrating the effect of 5 µM C77304 on PAC channel mutants of residues in the side portal region at their respective root pHs; voltage protocol as shown (n = 3 - 9). (C), Effects of 5 µM C77304 on the currents of PAC/R237A(at various pHs), PAC/E194A(at root pH) and PAC/F196A (at root pH) mutant channels, showing that the F196A and R237A mutations effectively eliminated the compound’s activating effect (n values as indicated in each bar). (D)-(E), Representative current traces (D) and concentration-response relationships (E) of C77304 inhibiting PAC-R237A/F196A mutant channel. The IC_50_s were calculated to be 5.7 ± 0.9 µM and 3.5 ± 0.9 µM at pH7.0 and pH6.4, respectively (n = 10 - 12).

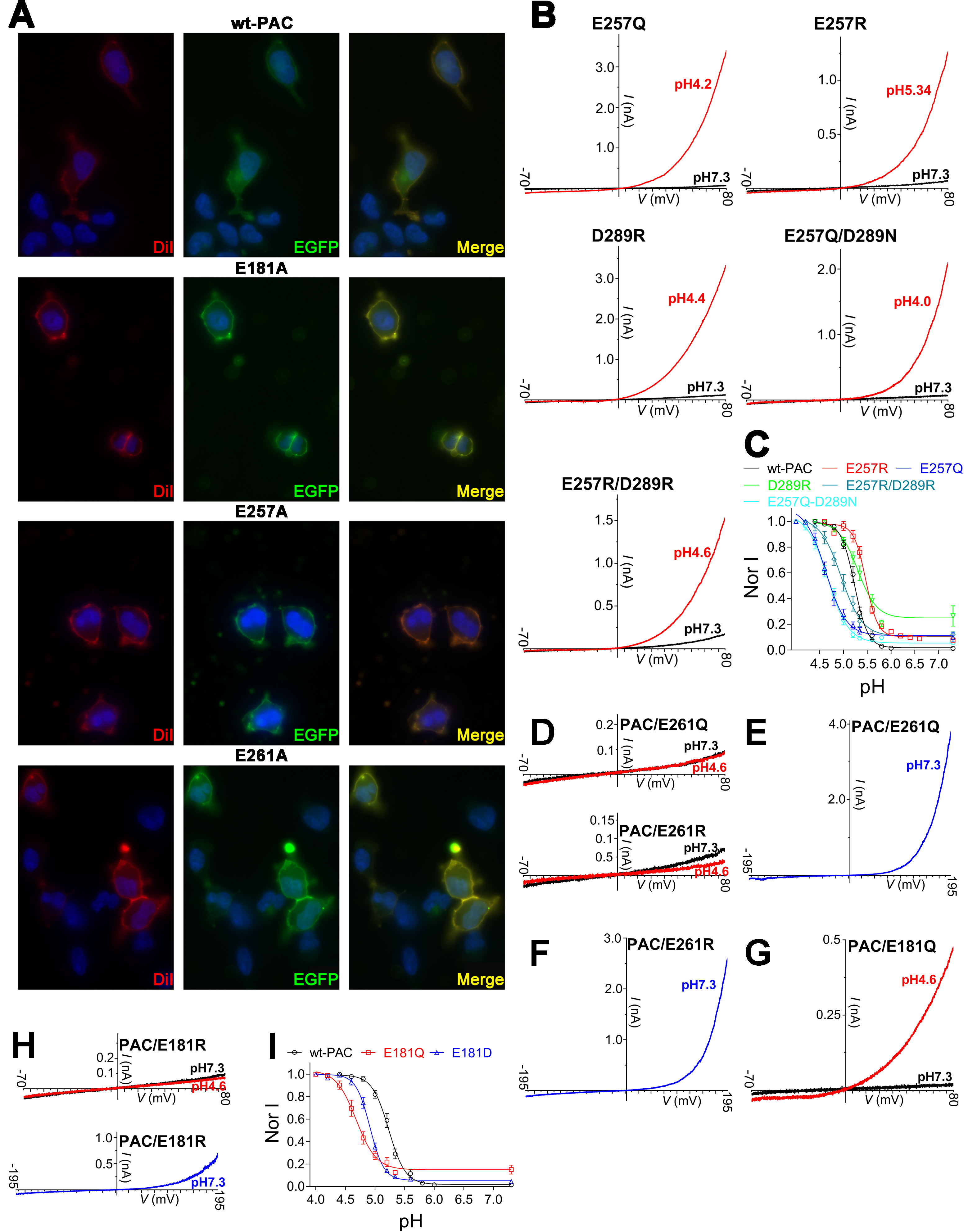

**Supplementary Fig. 6.** (A), Immunocytochemistry imaging revealed the membrane expression of GFP-tagged mutant channels (the green fluorescence), co-localized with the membrane marker DiI (the red fluorescence) Representative micrographs from three independent experiments are shown. (B), Representative current traces showing the PAC/E257Q, PAC/E257R, PAC/D289R, PAC-E257Q/D289N, and PAC-E257R/D289R mutant channels are functionally gated by protons (n = 5 – 9). (C), Current-pH relationships of PAC mutant channels as indicated, showing E257R, E257Q, E257Q/D289N, and E257R/D289R mutations remarkably shifted the curve to either acidic or alkaline directions, and D289R mutation caused a considerable basal opening at pH7.3 (n = 6 - 12). (D)-(F), Example current traces showing that the PAC/E261Q and PAC/E261R mutant channels were not functionally gated by proton (D) but by strong ramp depolarization at pH7.3 (E and F) (n = 12 - 31). (G), Typical proton-activated current of PAC/E181Q mutant channel (n = 7). (H), PAC/E181R mutant channel in response to acid (upper panel) and strong ramp depolarization at pH7.3 (lower panel) (n = 16 - 19). (I), Current-pH relationships of PAC/E181Q and PAC/E181D mutant channels compared with wt-PAC (n = 7 - 12).

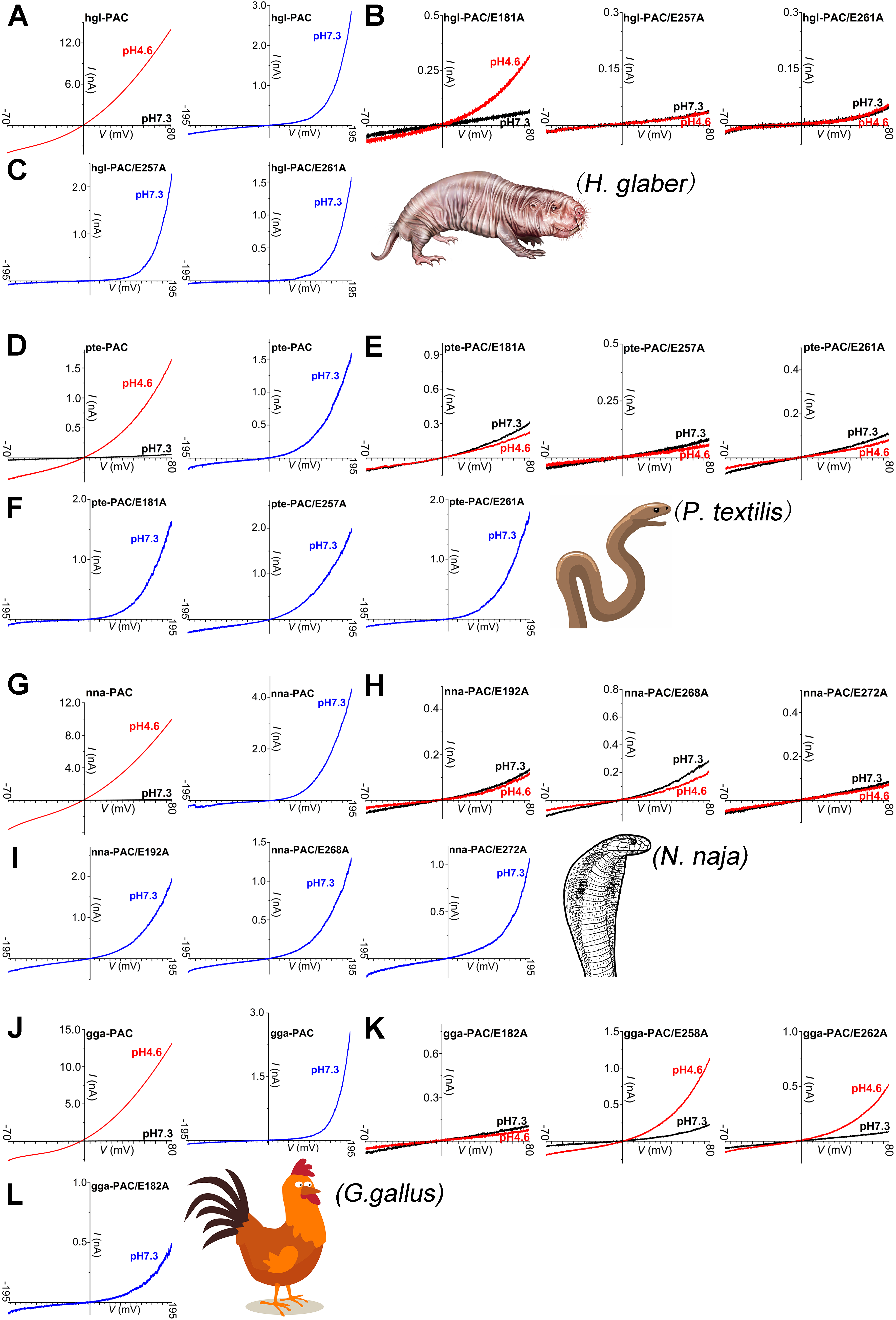

Supplementary Fig. 7 (continued)

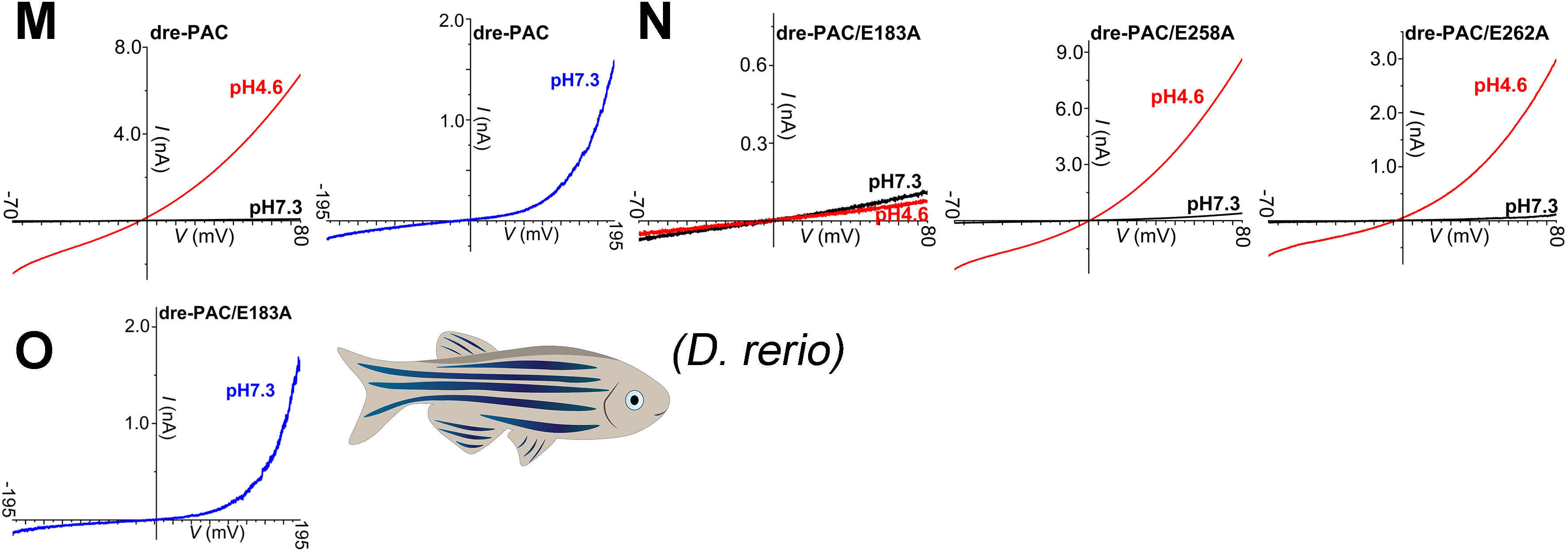

**Supplementary Fig. 7. Cross-species analysis of the proton sensing mechanism of PAC channel.** (A), Typical proton- or strong depolarization-activated currents of the PAC channel from *H. glaber (*hgl-PAC*)* (n = 11 - 20). (B), Mutating the human PAC E257 and E261 analogous sites (hgl-PAC/E257A and hgl-PAC/E261A) but not the E181 analogous site (hgl-PAC/E181A) in hgl-PAC eliminated its proton gating (n = 10 - 13). (C), Representative current traces showing hgl-PAC/E257A and hgl-PAC/E261A channels were activated by strong ramp depolarizations from -195 mV to +195 mV at pH7.3 (n = 17 - 19). (D), Representative proton- or strong depolarization-activated currents of the PAC channel from *P. textilis* (pte-PAC channel) (n = 8 - 13). (E), Example current traces showing mutating all the three residues analogous to the human PAC proton-sensing sites in pte-PAC (pte-PAC/E181A, pte-PAC/E257A, and pte-PAC/E261A) fully eliminated its proton gating (n = 7 - 10). (F), Representative strong depolarization activated currents of pte-PAC/E181A, pte-PAC/E257A, and pte-PAC/E261A mutant channels at pH7.3 (n = 10 - 13). (G), Typical proton- or strong depolarization evoked currents of the PAC channel from *N. naja* (nna-PAC channel) (n = 6 - 17). (H), Representative current traces showing E192A, E268A, and E272A mutations in nna-PAC (analogous to the human PAC E181, E257, and E261 site, respectively) abolished its proton gating (n = 7 - 12). (I), Typical strong depolarization activated currents of nna-PAC/E192A, nna-PAC/E268A, and nna-PAC/E272A mutant channels at pH7.3 (n = 14 - 18). (J), Typical proton- or strong depolarization-activated currents of the PAC channel from G. gallus (gga-PAC channel) (n = 7 - 14). (K), Example current traces showing E182A (analogous to human PAC E181) but not E258A and E262A (analogous to human PAC E257 and E261, respectively) mutations in gga-PAC eliminated its proton gating (n = 8 - 9). (L), Typical strong depolarization activated currents of gga-PAC/E182A mutant channel at pH7.3 (n = 13). (M), Typical proton- or strong depolarization-activated currents of the PAC channel from *D. rerio* (dre-PAC channel) (n = 12 - 23). (N), Representative current trace demonstrating that mutating residues analogous to human PAC E181 but not E257 and E261 in dre-PAC (dre-PAC/E183A but not dre-PAC/E258A and dre-PAC-E262A) fully eliminated its proton gating (n = 6 - 14). (O), Typical strong depolarization activated currents of dre-PAC/E183A mutant channel at pH7.3 (n = 9).

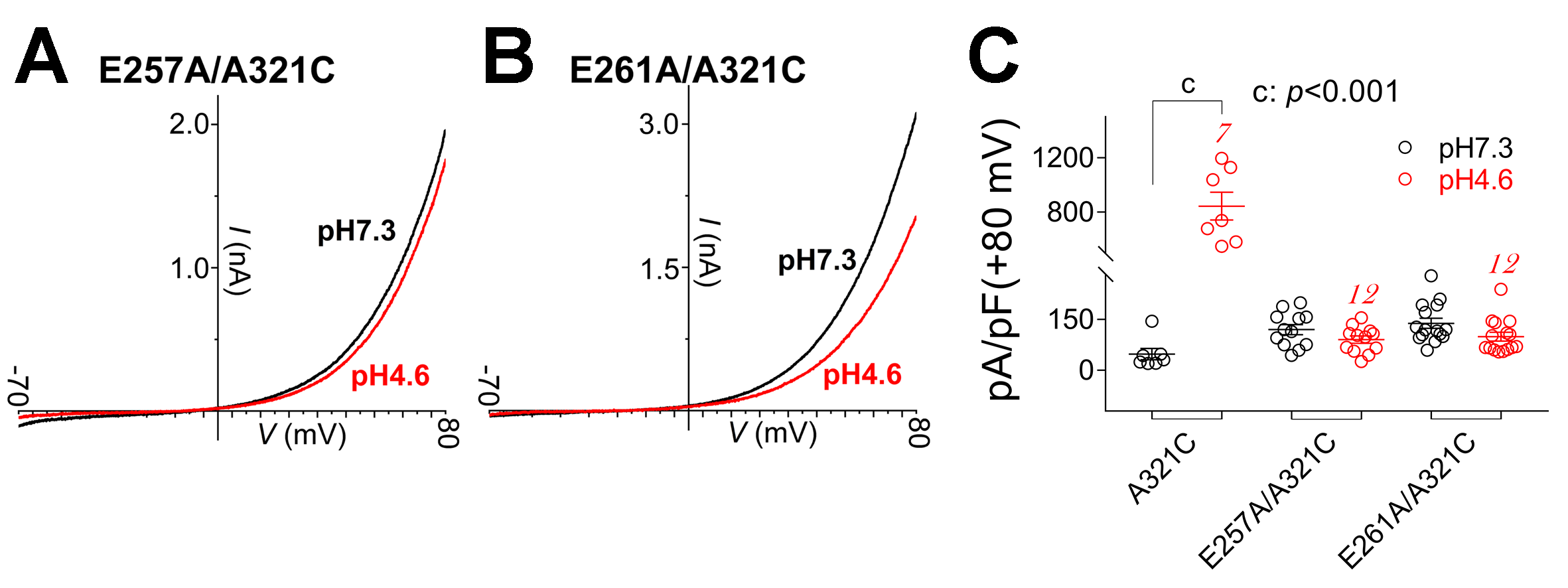

**Supplementary Fig. 8.** (A) and (B), Representative current traces demonstrating that the PAC-E257A/A321C (A) and PAC-E261A/A321C (B) double mutant channels are not activated by protons but exhibit considerable basal opening at pH7.3 (n = 12). Currents were recorded with a voltage ramp from -70 mV to +80 mV. (C), Summary analysis of the proton activated currents of PAC/A321C, PAC-E257A/A321C, and PAC-E261A/A321C mutant channels, significant differences between the pH7.3 and pH4.6 conditions were assessed using paired t-test; p value as indicated in the panel and n values as indicated in each bar.
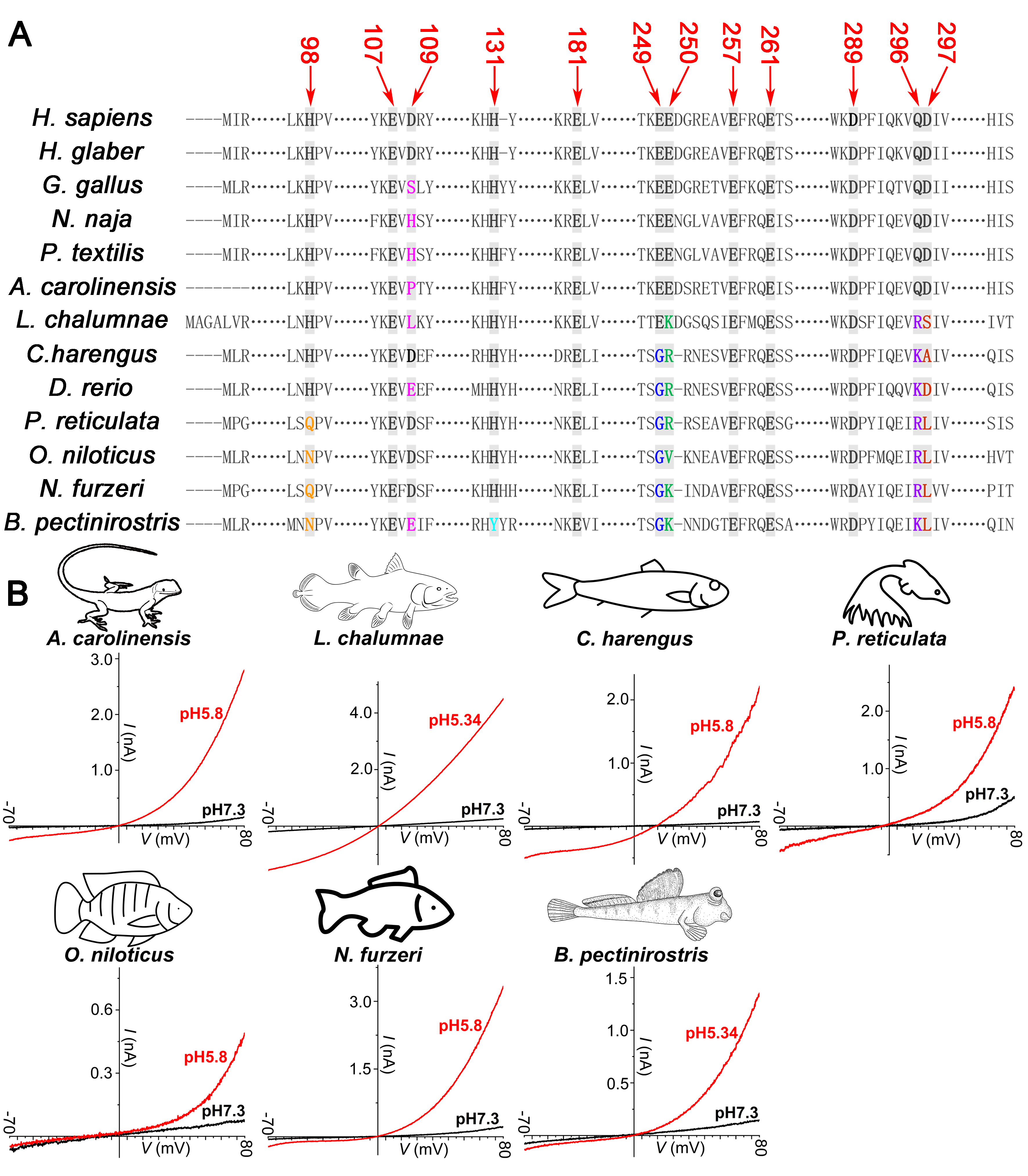

**Supplementary Fig. 9** (A), Sequence alignment of PAC channels from 13 different species as indicated, showing that residues E181, E257 and E261, but not the other gating-related residues are conserved across species (in human PAC numbering). In PAC channels from *P. reticulata*, *O. niloticus*, and *N. furzeri*, the proposed proton sensor H98 is mutated to glutamine or asparagine; in PAC channel from *B. pectinirostris*, the proposed proton sensors H98 and H131 are mutated to asparagine and tyrosine, respectively; in PAC channels from *C. harengus*, *P. reticulata*, *O. niloticus*, *N. furzeri*, and *B. pectinirostris*, the E249 and D297 analogous sites mutation are expected to abolish the proposed E107-E249 and E250-D297 carboxy-carboxylate interactions in the activated state; the D109, E250, and Q296 residues which were proposed to form the H98 binding pocket are also varied among orthologous PAC channels from lots of species. (B), Typical proton activated currents of PAC channels from *A. carolinensis*, *L. chalumnae*, *C. harengus*, *P. reticulata*, *O. niloticus*, *N. furzeri*, and *B. pectinirostris* (n = 4 – 9).

**Supplementary Table 1**

| **Channel** | **pH_50_** | **nH activation** | **BOP** | **n** |
| --- | --- | --- | --- | --- |
| wt-PAC | 5.23 ± 0.01 | -3.10 ± 0.42 | 0.01 ± 0.01^NS^ | 12 |
| wt-PAC/5 μM | 5.61 ± 0.01^d^ | -3.78 ± 0.50^NS^ | 0.04 ± 0.01^NS^ | 7 |
| wt-PAC/10 μM | 5.81 ± 0.03^d^ | -3.13 ± 0.65^NS^ | 0.04 ± 0.03^NS^ | 8 |
| M101A | 5.00 ± 0.04^d^ | -1.59 ± 0.20^NS^ | 0.001 ± 0.03^NS^ | 6 |
| V103A | 5.75 ± 0.02^d^ | -3.03 ± 0.50^NS^ | 0.01 ± 0.02^NS^ | 7 |
| Y105A | 5.42 ± 0.01^b^ | -3.35 ± 0.30^NS^ | -0.0002 ± 0.01^NS^ | 5 |
| H130A | 4.93 ± 0.01^d^ | -3.23 ± 0.25^NS^ | 0.004 ± 0.01^NS^ | 6 |
| Y132A | 5.07 ± 0.01^d^ | -3.18 ± 0.29^NS^ | 0.03 ± 0.01^NS^ | 9 |
| K178A | 5.41 ± 0.02^d^ | -3.50 ± 0.43^NS^ | 0.01 ± 0.01^NS^ | 8 |
| R180A | 5.06 ± 0.03^d^ | -3.11 ± 0.61^NS^ | 0.006 ± 0.03^NS^ | 7 |
| E181Q | 4.68 ± 0.03^d^ | -2.55 ± 0.50^NS^ | 0.14 ± 0.03^c^ | 8 |
| E181D | 4.91 ± 0.01^d^ | -3.68 ± 0.37^NS^ | 0.05 ± 0.01^NS^ | 7 |
| E194A | 5.32 ± 0.01^NS^ | -4.44 ± 0.55^NS^ | 0.03 ± 0.01^NS^ | 6 |
| F196A | 5.35 ± 0.14^NS^ | -1.88 ± 0.95^NS^ | 0.39 ± 0.06^d^ | 9 |
| D200A | 5.00 ± 0.008^d^ | -4.09 ± 0.28^NS^ | 0.009 ± 0.009^NS^ | 7 |
| E209A | 5.07 ± 0.02^d^ | -3.38 ± 0.56^NS^ | 0.01 ± 0.02^NS^ | 8 |
| E224A | 5.00 ± 0.01^d^ | -3.40 ± 0.33^NS^ | 0.01 ± 0.01^NS^ | 6 |
| S228A | 4.14 ± 0.02^d^ | -2.92 ± 0.46^NS^ | 0.02 ± 0.02^NS^ | 7 |
| S229A | 5.13 ± 0.01^a^ | -3.27 ± 0.23^NS^ | 0.01 ± 0.01^NS^ | 6 |
| W230A | 5.53 ± 0.03^d^ | -2.71 ± 0.44^NS^ | -0.003 ± 0.03^NS^ | 6 |
| K231A | 6.25 ± 0.03^d^ | -2.18 ± 0.31^NS^ | 0.004 ± 0.03^NS^ | 9 |
| R237A | 5.87 ± 0.06^d^ | -1.18 ± 0.21^b^ | 0.005 ± 0.05^NS^ | 6 |
| W239A | 4.62 ± 0.06^d^ | -1.71 ± 0.31^NS^ | 0.08 ± 0.03^NS^ | 5 |
| K241A | 4.90 ± 0.01^d^ | -3.05 ± 0.33^NS^ | 0.02 ± 0.01^NS^ | 7 |
| E249A | 4.93 ± 0.01^d^ | -4.06 ± 0.45^NS^ | 0.01 ± 0.01^NS^ | 6 |
| E250A | 5.01 ± 0.01^d^ | -3.50 ± 0.29^NS^ | 0.02 ± 0.01^NS^ | 6 |
| D251A | 5.07 ± 0.01^d^ | -3.18 ± 0.31^NS^ | 0.01 ± 0.01^NS^ | 8 |
| E257Q | 4.62 ± 0.05^d^ | -2.02 ± 0.37^NS^ | 0.11 ± 0.03^NS^ | 6 |
| E257R | 5.46 ± 0.01^d^ | -3.46 ± 0.36^NS^ | 0.10 ± 0.01^NS^ | 7 |
| R259A | 4.49 ± 0.05^d^ | -2.10 ± 0.40^NS^ | 0.03 ± 0.04^NS^ | 7 |
| Q260A | 4.83 ± 0.03^d^ | -2.71 ± 0.46^NS^ | 0.0002 ± 0.03^NS^ | 8 |
| E286A | 4.78 ± 0.02^d^ | -2.94 ± 0.42^NS^ | 0.002 ± 0.02^NS^ | 5 |
| K288A | 4.48 ± 0.02^d^ | -3.53 ± 0.52^NS^ | 0.03 ± 0.02^NS^ | 5 |
| D289A | 4.29 ± 0.04^d^ | -1.80 ± 0.26^NS^ | 0.04 ± 0.02^NS^ | 7 |
| D289R | 5.29 ± 0.05^NS^ | -2.17 ± 0.52^NS^ | 0.24 ± 0.04^d^ | 8 |
| Q293A | 5.26 ± 0.007^NS^ | -3.93 ± 0.27^NS^ | 0.007 ± 0.008^NS^ | 6 |
| V295A | 5.54 ± 0.02^d^ | -2.09 ± 0.20^NS^ | -0.005 ± 0.02^NS^ | 6 |
| E257Q/D289N | 4.66 ± 0.03^d^ | -1.98 ± 0.25^NS^ | 0.05 ± 0.02^NS^ | 7 |
| E257R/D289R | 4.95 ± 0.06^d^ | -1.94 ± 0.40^NS^ | 0.10 ± 0.04^NS^ | 6 |

Supplementary Table 1: Summary of proton gating parameters of wild-type (wt) and mutant PAC channels tested in this study. Abbreviations: pH_50_, half activation pH; nH activation, slope factor of pH-current relationship; BOP, basal opening proportion at neutral pHs as determined by fitting the pH-current relationships; n, number of separate experimental cells; a, *p*<0.05; b, *p*<0.01; c, *p*<0.001; d, *p*<0.0001; Statistical differences were assessed by ONE-WAY ANOVA with post-hoc analysis using the Dunnett method (wt-PAC as control). Data are presented as MEAN ± SEM.
